# Regional climate affects salmon lice dynamics, stage structure, and management

**DOI:** 10.1101/574830

**Authors:** Amy Hurford, Xiunan Wang, Xiao-Qiang Zhao

## Abstract

Regional variation in climate can generate differences in population dynamics and stage structure. Where regional differences exist, the best approach to pest management may be region specific. Salmon lice are a stage structured marine copepod that parasitizes salmonids at aquaculture sites worldwide, and have fecundity, development, and mortality rates that depend on temperature and salinity. We show that in Atlantic Canada and Norway, where the oceans are relatively cold, salmon lice abundance decreases during the winter months, but ultimately increases from year-to-year, while in Ireland and Chile, where the oceans are warmer, the population size grows monotonically without any seasonal declines. In colder regions, during the winter the stage structure is dominated by the adult stage, which is in contrast to warmer regions where all stages are abundant year round. These differences translate into region specific recommendations for management: regions with slower population growth have lower critical stocking densities, and regions with cold winters have a seasonal dependence in the timing of follow-up chemotherapeutic treatments. Predictions of our salmon lice model agree with empirical data, and our approach provides a method to understand the effects of regional differences in climate on salmon lice dynamics and management.

## 1. Introduction

Population dynamics vary regionally and best approaches to management for pest species, such as salmon lice (*Lepeophtheirus salmonis*), are likely region specific (Robbins et al., 2010). Regional variation in population dynamics are illustrated in Fennoscandian microtine voles (*Microtus* sp.) where southerly populations are non-cyclic, while more northerly populations have multi-annual cycles (Hanski et al., 1991). In addition, for cyclic populations of microtine voles, gypsy moth (*Lymantria dispar (L.)*), and jack pine budworm (*Choristoneura pinus*)), cycle lengths (Bjørnstad et al., 1995, 2010), and amplitudes (Bjørnstad et al., 1995) vary by region. In insects, the number of generations per year is referred to as voltinism, and multivoltinism occurs in insect species found in regions that are warm (Beck and Apple, 1961) or have warmed (Altermatt, 2010). In pest species, these regional differences may suggest different best approaches to management and control, for example, the optimal timing of mosquito fogging to reduce the impacts of dengue fever depends on the timing of the onset of the wet season (Oki et al., 2011), which may vary between regions.

While a range of factors underlie regional differences in population dynamics (Bjørnstad et al., 1995, 2010; Hanski et al., 1991), climate varies regionally and for many species there is a direct relationship between environmental variables and life history characteristics, including the timing of oviposition and voltinism (Powell and Logan, 2005; Culos and Tyson, 2014), which will affect population dynamics. Recent studies have parameterized the temperature dependencies of fecundity, mortality, and maturation rates to predict the population dynamics of tea tortix (*Adoxophyes honmai*; Nelson et al. 2013), mosquitoes (*Culex pipiens*; Ewing et al. 2016), biting midges (*Culicoides* sp.; White et al. 2017), and salmon lice (Rittenhouse et al., 2016). While ecophysiological models that utilize laboratory data to parameterize models of natural population dynamics have been criticized (Davis et al., 1998), ultimately these models have performed well for many species (Hodkinson, 2001; Nelson et al., 2013), generate testable predictions, and identify the most critical areas for future model improvement.

Salmon lice have distinct life stages distinguished by different fecundity, maturation, and mortality rates, and mathematical models accounting for stage structure, together with density dependence, exhibit a range of complex population dynamics including multigenerational cycles and chaos (Gurney et al., 1980). For stage structured populations, the relationships between temperature and fecundity, maturation, and mortality rates are potentially different for each life stage. Insect diapause is an illustrative example: diapause is a developmental delay to facilitate species survival in adverse environmental conditions, but usually occurs in only one life stage (Chapman, 2012). For stage structured models, we may consider the dynamics of the stages relative to each other, which may include discrete non-overlapping generations (Gurney et al., 1980), and ‘holes’ in the population structure, where one or more stages are absent for a length of time (de Roos et al., 2009). Pest management strategies often target a specific life stage, and so understanding stage structured population dynamics is a relevant consideration for effective pest management.

Salmon aquaculture can occur in regions where large differences in sea temperature occur between seasons. The consequences of seasonal variation in life history parameters may be simple annual cycles or more complex multiyear dynamics depending on the strength of seasonality and the parameters affected (Altizer et al., 2006). Mathematical models that consider seasonality can be formulated as dynamical systems with periodic coefficients, or delay differential equations when a threshold level of development is necessary for maturation to the next life stage (Nisbet and Gurney, 1983). Recently, theory has been developed to calculate threshold quantities such as the basic reproductive ratio, *R*_0_, for these more complex dynamical systems (Diekmann et al., 1990; Bacaer and Guernaoui, 2006; Zhao, 2017). The basic reproductive ratio characterizes the stability of an extinction equilibrium, can be interpreted as the lifetime reproduction of an average individual, and can be reframed as a threshold host density characterizing outbreak dynamics (Frazer et al., 2012). Mathematical models that ignore seasonality are easier to analyze, but may yield erroneous results (Mitchell and Kribs, 2017) and provide insufficient information to inform the best timing of treatments within a season.

We study salmon lice because the relationship between the environment and life history parameters, and the salmon lice population dynamics themselves, have been well studied in the past (Tucker et al., 2002; Stein et al., 2005; Rittenhouse et al., 2016; Groner et al., 2016; Frazer et al., 2012). In addition, salmon lice have a substantial adverse economic impact on salmon farms (Costello, 2009). We consider seasonal patterns in sea surface temperature and salinity and develop the mathematical theory to calculate the basic reproductive ratio for the system of delay differential equations that we use to model salmon lice dynamics. We consider 11 salmon farming regions, and find that in Atlantic Canada and Norway, where winter oceans are colder: (1) the adult salmon lice population will decrease during the winter, and few other life stages will be present; and (2) the timing of follow-up chemotherapeutic treatments, to affect a second generation parasites, will vary substantially depending on the time of year. Both these results suggest that regional differences in population dynamics translate into regional differences in optimal strategies for pest management on salmon farms.

## 2. Methods

### 2.1 Model formulation

To investigate the population dynamics of salmon lice, we use the model derived in Rittenhouse et al. (2016), which considers seasonal environments, while still being simple enough to calculate critical stocking densities and life stage durations. Salmon lice are assumed to be in one of four stages: nauplii, *P* (*t*), copepodids, *I*(*t*), chalimi and pre-adults, *C*(*t*), and adult females, *A*(*t*). The nauplius stage includes all planktonic non-infectious stages, and collectively nauplii and copepodids are referred to as ‘larvae’, with the abundance of larvae calculated as *P* (*t*) + *I*(*t*). Adult males are not explicitly considered in the model formulation, but are assumed to be equal in abundance to adult females. The chalimus/pre-adult, adult female, and adult male stages are collectively referred to as ‘parasites’ and the abundance of the parasite stages is calculated as *C*(*t*) + 2*A*(*t*). The equations for the stage structured dynamics of salmon lice are,

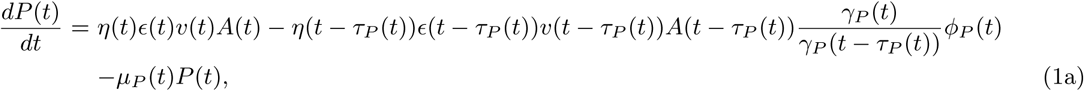

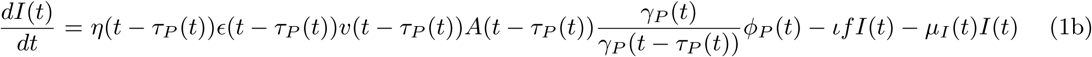

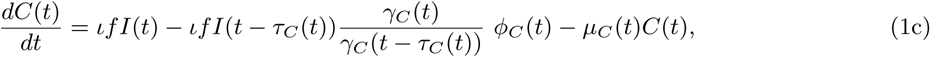

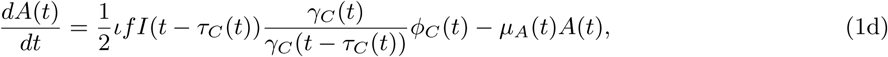

where,

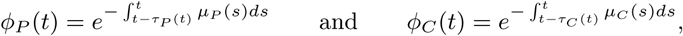

and where the lengths of the maturation delays are calculated by solving the equations,

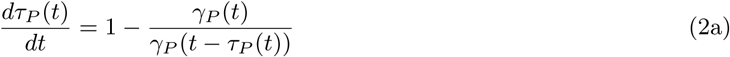

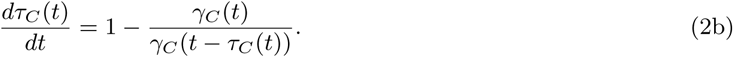

The terms in each delay differential equation in system (1) correspond to the rate of entry into a stage via the hatching of eggs or maturation, the rate of loss due to maturation or attachment (except for in the adult female stage because this is the terminal stage of development), and the rate of mortality, and appear in that order. For complete details of the model derivation see Section S.2 of the Supplementary Online Material (SOM), and for definitions of all parameters see Table S.1. The delay equations (2) assume that a cumulative amount of temperature-dependent development must be completed for maturation to the copepodid and adult female stages. These functions for the time delays (equations (2)), and the other time-dependent parameters (i.e., the number of eggs per clutch, *η*(*t*)), are ultimately functions of sea surface temperature, *T* (*t*), and/or salinity, *S*(*t*), which are seasonal and have a period, *ω*, of one year.

The model (equations (1) and (2)) does not consider any negative density dependence because on salmon farms interventions are used to reduce the number of salmon lice before their densities are large enough to decrease the per capita fecundity and survival rates. For example, interventions may occur when the density of lice is 1-2 adult females per fish (Krkosek et al., 2010), but densities of >20 adult females per fish are recorded (for example, see the dataset provided in Marty et al. 2010), suggesting that interventions occur when densities are relatively low. The model (equations (1) and (2)) does not explicitly consider inflow and outflow of larval stages from the salmon netpens due to ocean currents, but this flow may be understood as implicitly included in the parameterization of the mortality rate of nauplii, *µ*_*P*_ (*t*), and the copepodid attachment rate, *ι*, as discussed in Section S.1 of the SOM.

The salmon farming sites we consider are: Region X, Chile (CH); 2 sites west of Ireland (IM3, IM4); 3 sites in British Columbia, Canada (Broughton Archipelago, BCB, Central Coast, BCC, and Vancouver Island, BCV); 3 sites in Atlantic Canada (New Brunswick, NB, Nova Scotia, NS, and Newfoundland, NL), and 2 sites in Norway (Ingøy, NIN, and Lista, NLI) (Figure 1). Temperature and salinity are assumed to be periodic or constant functions (based on fit) and locations without salinity data are assumed to have a constant salinity of 31 psu (equal to the mean salinity from sites with reported salinity, but excluding NL which had very low salinity). To facilitate comparisons with the Northern Hemisphere sites, the temperate data from Region X, Chile is shifted by 182.5 days.

**Fig. 1.**
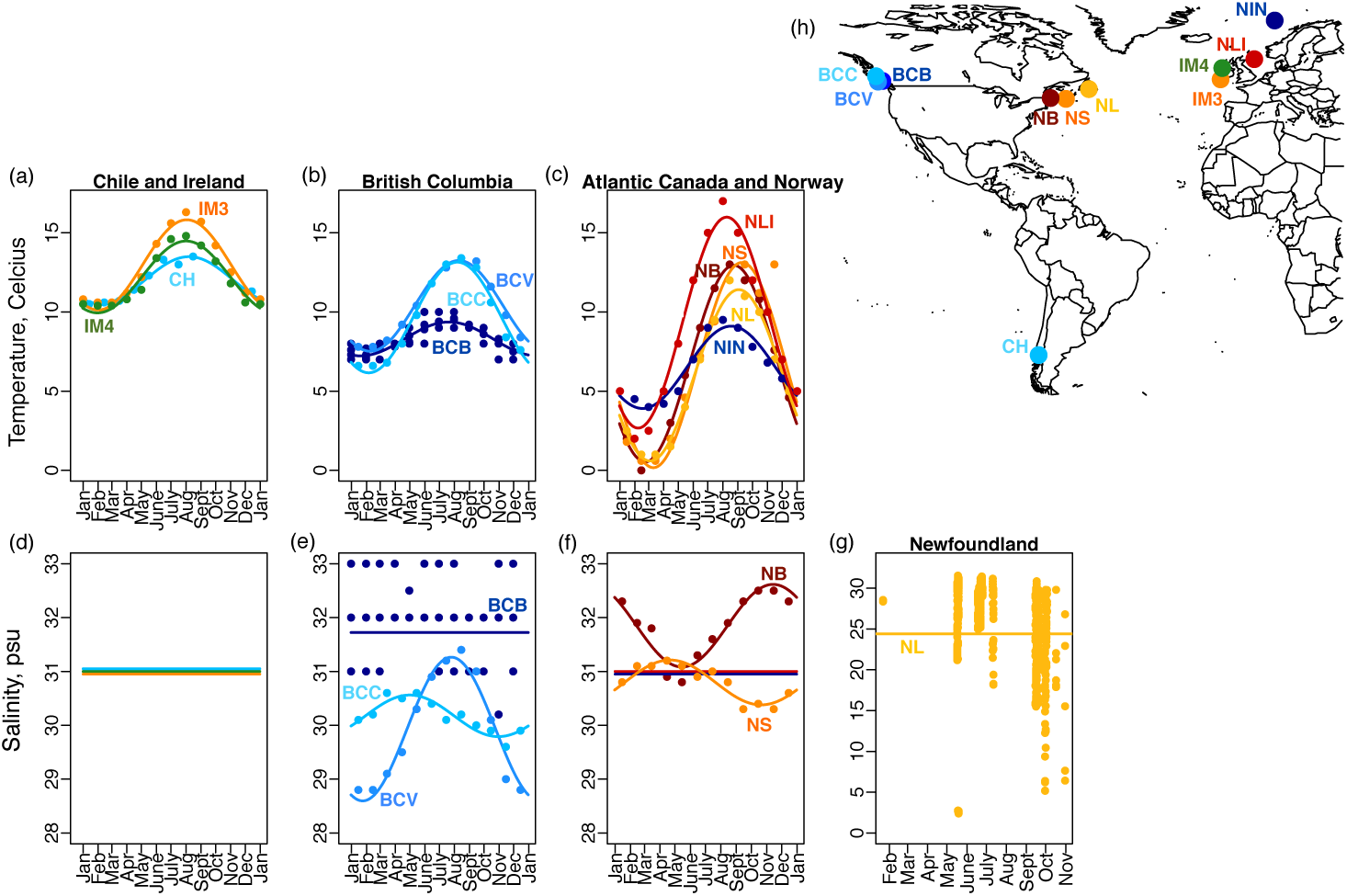
Environmental data for 11 salmon farming sites and their locations. Sea surface temperature (a-c) and salinity data (d-g) for salmon farming sites (h) grouped by minimum temperature. (a) Sites in Chile (sky blue) and Ireland (IM3 - orange, and IM4 - green) have the highest yearly minimum temperatures. Salinity data is not available for CH, IM3 and IM4, so a constant salinity of 31 psu is assumed (d). (b) Sites in British Columbia, Canada (BCB - dark blue, BCV - mid blue, and BCC - sky blue) have intermediate sea surface temperatures (b), with salinity shown in (e). (c) Sites in Atlantic Canada (NS- orange, NB - dark red, NL - yellow) and Norway (NIN - red, NLI - dark blue) have the coldest minimum temperatures with salinity shown in (f) and (g). Salinity data was not available for the Norway sites and was assumed to be 31 psu. (g) Surface salinity for the Newfoundland site is shown in a separate panel because here salinity was much lower than any of the other sites. Additional details are provided in Section S.1.

Data from laboratory experiments for *Lepeophtheirus salmonis* are used to estimate life history parameters that are functions of temperature and salinity. The density of fish at a site, *f*, and the copepodid attachment rate, *ι*, were estimated using data from the BCB site. Aside from the functions for temperature, *T* (*t*), and salinity, *S*(*t*), (Table S.3) all other parameters (Table S.1) are the same across sites, so that we can isolate the effect of local temperature and salinity trends on salmon lice population dynamics.. Complete details describing the model parameterization are provided in Section S.1.

### 2.2 Model analysis

Motivated by the Floquet theory of periodic ordinary differential equations and Lemma 1 in Wang and Zhao (2017), we refer to 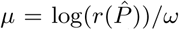 as the principal (or leading) Floquet exponent, where *ω* = 1 year is the length of the periodicity and 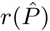 is the spectral radius of the Poincaré map, 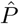, corresponding to system (S.21), which is (1) without the *dP* (*t*)*/dt* and *dC*(*t*)*/dt* equations (since in Section S.3, we show that the dynamics of system (1) are determined by the dynamics of (S.21)). In Section S.3, we define the basic reproductive ratio, *R*_0_, the number of second generation adult females produced by an average adult female during its lifespan, and we prove the following analytic result:

#### Theorem 1

*System* (1) *and* (2) *admit the following threshold dynamics:*

*(i) If R*_0_ *<* 1 ⇔ *µ <* 0, *then* (0, 0, 0, 0) *is globally attractive;*
*(ii) If R*_0_ > 1 ⇔ *µ* > 0, *then the nontrivial solutions go to infinity as t → ∞.*

We define the critical stocking density, *f*_*crit*_, as the number of fish on a farm such that *f < f*_*crit*_ implies *R*_0_ *<* 1, and hence, salmon lice cannot persist. We numerically solve the system of equations (1) and (2) using the `PBSddesolve` (Couture-Beil et al., 2016) package for *R*. We investigate the effects of ignoring seasonality in sea surface temperature and salinity by setting *T* (*t*) = *a* and *S*(*t*) = *c* and *ι* = 2.4 *×* 10^−9^ (the fitted value when seasonality was ignored).

### 2.3 The follow-up treatment window

For chemotherapeutic treatments, such as cypermethrin, only the parasitic salmon lice stages are affected. The follow-up treatment window is defined as a range of days, *T*_1_ to *T*_2_, after an initial treatment such that: (a) all larval lice that escaped the initial treatment will be in the parasite stage when the follow-up treatment occurs; and (b) none of the larval lice that escaped the initial treatment have yet reached the reproductive adult female stage. Since the development level of nauplii can never decrease (i.e., *γ*_*P*_ (*t*) is non-negative) the beginning of the treatment window, *T*_1_, so as to satisfy (a) can be calculated as the time it takes the least mature larvae at the time of the initial treatment (i.e., a recently hatched nauplius) to reach the chalimus stage. The time a larval louse spends in the copepodid stage depends on its ability to find a salmonid host, which is dependent on oceanographic currents that our model does not consider, so we assume that the time spent in the copepodid stage is 10 days as reported in Tucker et al. (2002). For a given current temperature of *a* degrees Celcius, and making the simplifying assumption that past temperatures are equal to the current temperature, we calculate the start of the treatment window as,

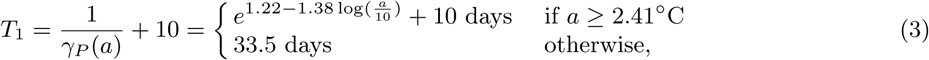

which is based on the parameterization of *γ*_*P*_ (*T*) from laboratory experiments (Section S.1). Similarly, since the development level of chalimii/pre-adults cannot decrease (*γ*_*C*_ (*T*) *≥* 0), the end of the treatment window, *T*_2_, so as to satisfy (b) can be calculated as the time it takes for the most mature larvae (a copepodid that attaches to a salmon, thus entering the *C*(*t*) stage immediately after the initial treatment) to mature to the adult female stage. As such, for a given current temperature of *a* degrees Celcius, and using the simplifying assumption that past temperatures are equal to the current temperature, we calculate the end of the treatment window as,

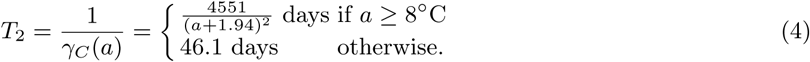

The existence of a follow-up treatment window (*T*_2_ > *T*_1_) is not guaranteed, however, for the parameterization of this salmon lice model (specifically, the functions for *γ*_*P*_ (*T*) and *γ*_*C*_ (*T*)), the follow-up treatment window exists for sea surface temperatures less than 17.9°C (i.e., by rearranging the inequality *T*_2_ > *T*_1_ to isolate *a*, see (3) and (4)) and we note that sea surface temperature never exceeds this temperature at any of our sites (Figure 1).

To calculate the follow-up treatment window when the simplifying assumption is relaxed to allow for seasonality, whereby past temperatures can be warmer or cooler than the current temperature, we calculate the values of *τ*_*P*_ (*t*) and *τ*_*C*_ (*t*), which are the lengths of time spent in the nauplius and chalimus/pre-adult stages for a salmon louse that exits these stages at time *t*. We numerically solve equations (2), and we again assume that the time spent in the copepodid stage is 10 days (see Section S.5 in the SOM). Complete details of our numerical methods are provided in Section S.4 and S.5, and all code to produce our results is publically available at Hurford et al. (2019).

## 3. Results

Floquet exponents, *µ*, are positive and *R*_0_ is greater than one at all sites except for Newfoundland (NL) indicating that the salmon lice extinction equilibrium is unstable and the number of salmon lice increases from year-to-year (Theorem 1; Figure 2a). At the Newfoundland site, the extinction equilibrium is stable and the number of salmon lice decreases from year-to-year. To determine whether the stability of the extinction equilibrium at the Newfoundland site is due to low salinity, we assume constant salinity of 31 psu and denote this site as (NL). With the elevated salinity (31 psu) the Floquet exponent at the (NL) site is positive and *R*_0_ is greater than 1. We calculated the critical stocking density at each site as ranging between 61,406 (IM3) and 309,770 (NIN) fish per farm (Figure 2b), and these critical stocking densities were closely related to the Floquet exponent (Figure 2d).

**Fig. 2.**
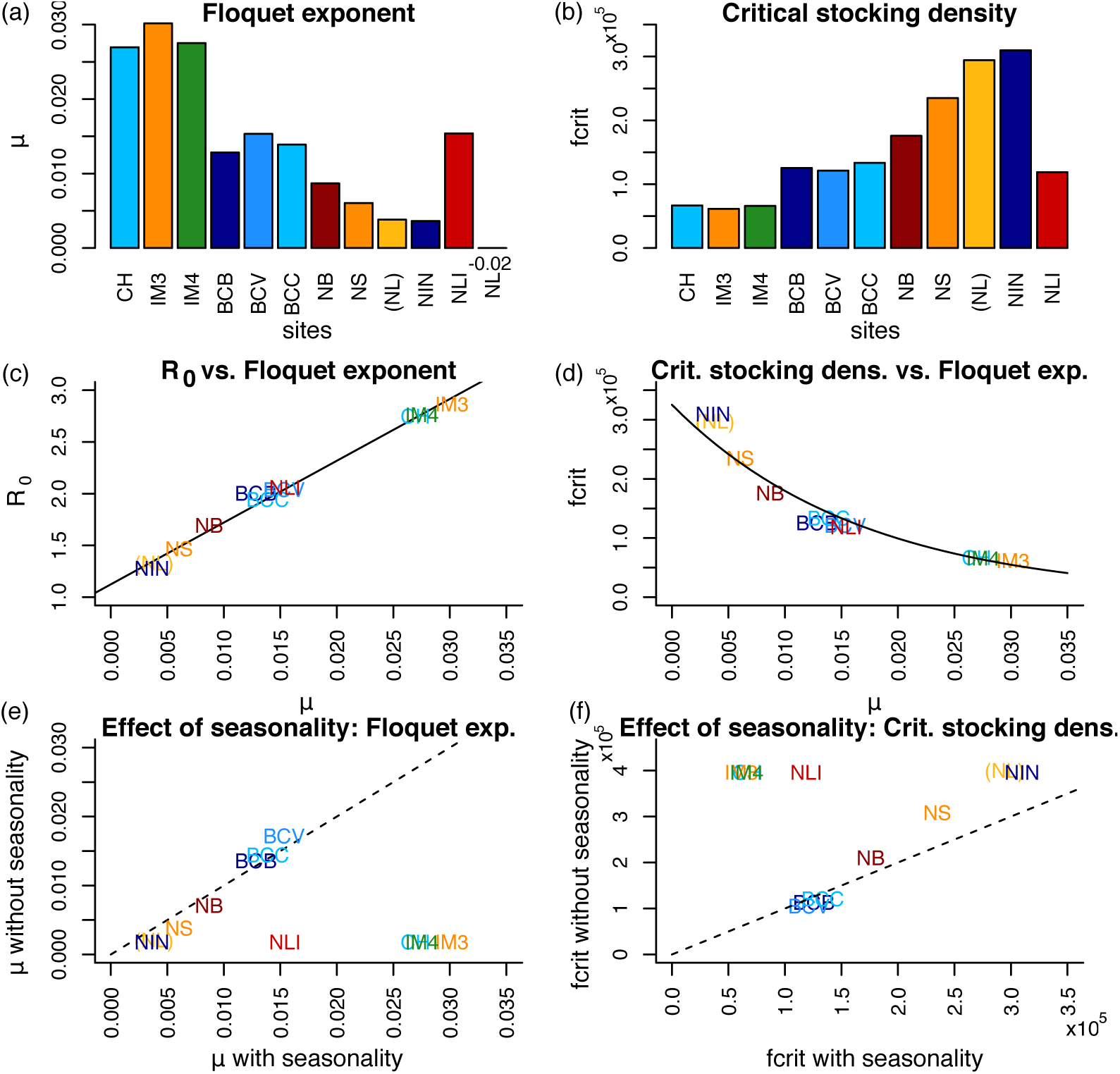
*R*_0_ is strongly correlated with the leading Floquet exponent, *µ*, critical stocking densities are higher in Atlantic Canada, and ignoring environmental periodicity can result in large overestimates of the critical stocking densities. (a) The Floquet exponents are all positive except for Newfoundland (NL, *µ* = − 0.02). For all other sites, including Newfoundland with salinity at 31 psu (NL), the Floquet exponent is positive indicating an increase in the abundance of salmon lice from year-to-year. (b) Critical stocking densities range between 61,405 (IM3) and 309,770 (NIN) fish per farm. (c) The basic reproductive ratio and the leading Floquet exponent are linearly related with the best fit line *R*_0_ = 1.12 + 59.7*µ* (solid line), *R*^2^ = 0.99. (d) The Floquet exponent is closely related to the critical stocking density *f*_*crit*_ = exp(1.18 − 59.4*µ*) (*R*^2^ = 0.96). Ignoring the periodic variation in sea surface temperature and salinity results in an underestimate of *µ* (e), and an overestimate of *f*_*crit*_ (f), particularly at the CH, IM4, IM4 and NLI sites. The dashed line shows equal values when seasonality is ignored and included.

For the 11 salmon farming sites we considered, we found a strong linear relationship between the Floquet exponent, *µ*, and the basic reproductive ratio, *R*_0_ (Figure 2c, *R*^2^ = 0.99). We found that ignoring seasonality can substantially underestimate the Floquet exponent (Figure 2e) and overestimate the critical stocking density (Figure 2f). These inaccuracies were most pronounced for the sites with the warmest summer temperatures (CH, IM3, IM4 and NLI). At one site off the coast of Ireland (IM3), when seasonality is ignored, the critical stocking density is estimated as 396,088 fish, which is more than six times the value when seasonality is considered.

We find that in different locations salmon lice have qualitatively different dynamics and different stage structures (Figure 3). The sites in Atlantic Canada (NB and NS) have much slower population growth rates (*µ <* 0.01 and *R*_0_ *<* 1.8; Figure 3a), and in addition, the cold winter temperatures give rise to declines in the abundance of all stages (Figure 3b,c,d), although these abundances will rebound when environmental conditions become more favourable since *R*_0_ > 1 (Figure 2c). The slower population growth for the sites in Atlantic Canada is most likely due to colder winter temperatures (Figure 1) resulting in longer generation times (Figure 4).

**Fig. 3.**
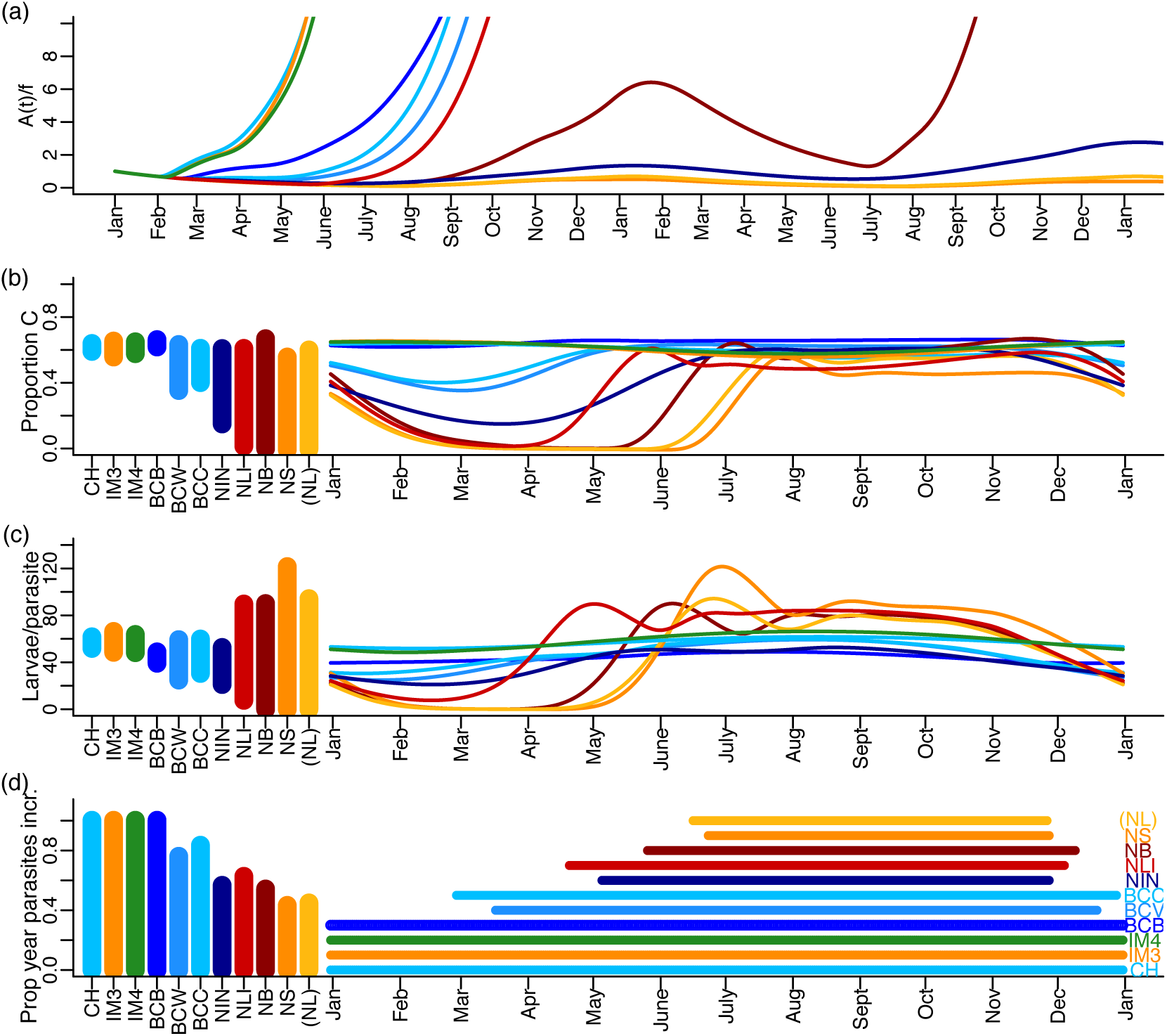
Stage structured dynamics for salmon lice at 11 sites. (a) The number of adult female salmon lice per fish increases much more rapidly at the warmer sites (CH - sky blue, IM3 - orange, IM4 - green) than at the colder sites (NB - dark red, NS - orange, (NL) - yellow and NIN - dark blue). (b) At the warmer sites, the proportion of parasites that are chalimi is relatively constant 0.55-0.66, but for the sites that are colder, the proportion of parasites that are chalimi/pre-adults can drop to near zero; such temperatures occur during the winter in northern Norway (NLI) and Atlantic Canada (NB, NS and (NL)). In the winter at these same sites, the number of larvae per parasite drops to near zero, while at sites with warmer winter temperatures this number ranges between 48 and 66 larvae per parasite (c). (d) At the warmer sites the number of parasites per fish increases year round: population growth is monotonic, but at sites with prolonged cold periods, the number of parasites per fish can decrease for up to 57% of the year (NS). In (a) the vertical axes does not extend further because our model assumes no density dependence and is only intended to apply to low densities.

**Fig. 4.**
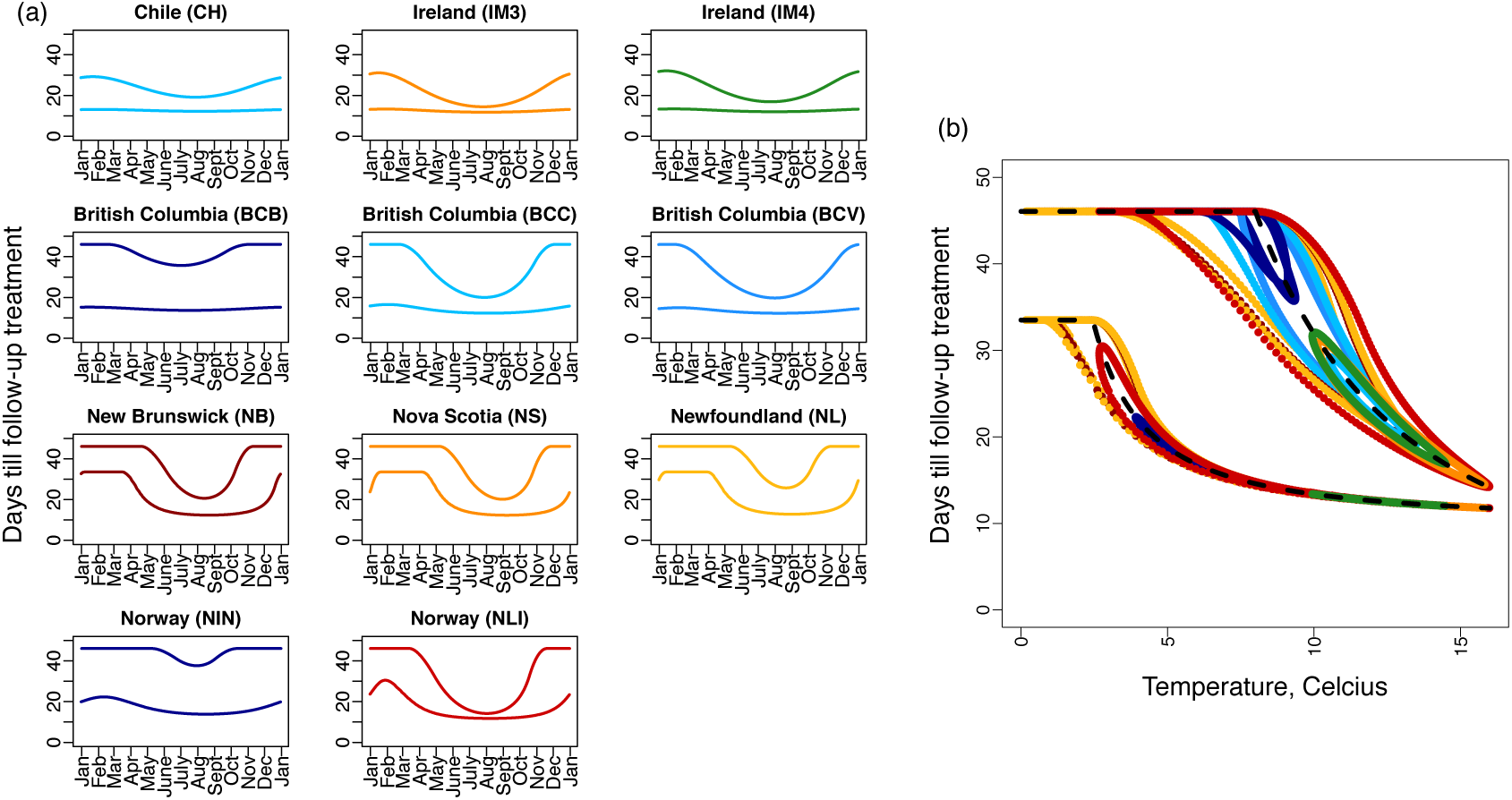
The timing of follow-up treatments vary seasonally and regionally. Chemotherapeutic treatments such as cypermethrin affect only the parasite stages of salmon lice. The follow-up treatment window is defined as: (i) beginning at *T*_1_ when a nauplius that was newly hatched at the initial treatment has matured to the chalimus stage, and (ii) ending at *T*_2_ when a copepodid that attached to a salmon immediately after the initial treatment has matured to the adult female stage. (a) Follow-up treatment windows for the 11 different salmon farming sites. (b) An approximation for the follow-up treatment window that assumes past and current temperatures are equal (dashed black line; equations (3) and (4)) matches well with the follow-up treatment windows using the seasonal delay equations (equations (2); shown in colours corresponding to each of the 11 salmon farming sites) when temperatures are not strongly seasonal.

Figure 3b and c reveals that the stage structure of the salmon lice population may vary substantially between regions and within a season. At the sites with warmer winter temperatures (CH, IM3, IM4), the stage structure is relatively unchanging year round: for every parasite there is 48-66 larvae and 55-66% of the parasite population is chalimi/pre-adults. The Atlantic Canada sites with cold winter temperatures (NB, NS and (NL)) demonstrate holes in the population structure where during the winter months adult salmon lice dominate the population structure. While in Atlantic Canada there may be nearly 1 larvae for every 3 parasites during the winter, when compared to larval abundance during the summer months (>60 larvae per parasite), larvae are near absent. Similar, but less extreme, changes in the population structure are observed for the sites in Norway (NIN and NLI), and the structure at the BC sites (BCB, BCC, and BCV) are intermediate, but more similar to the warmer sites in Chile and Ireland (Figure 3b,c, CH, IM3, and IM4). The stage structure predicted for the Broughton Archipelago site is compared to observations in Table 1.

**Table 1.** Model validation: a comparison of observations from the Broughton Archipelago with our model predictions.

| Quantity | Observation | Prediction | Details |
| --- | --- | --- | --- |
| Critical stocking density of the Broughton Archipelago | 17.3 kT<br>Frazer et al. (2012) | 16.3 kT | The critical stocking density for BCB is 125,517 fish (Figure 2b). Prediction assumes 5 kg/fish (Byrne et al., 2008) and 26 farms in the Archipelago (Marty et al., 2010). |
| Nauplii per copepodid | $87.8/5.2 = 16.9$<br>Byrne et al. (2008) | 0.8 | |
| Larvae per parasite | 8.3<br>Byrne et al. (2008) | 39-49 | We estimate the slope of the regression in Figure 5 of Byrne et al. (2008) as 0.25. Byrne et al. (2008) assumes 0.15 kg/m<sup>3</sup> and 5 kg/fish. |
| Percentage of parasites that are chalimi and pre-adults | 51<br>Marty et al. (2010) | 62-66 | Prediction calculated from a subset of the data provided in Marty et al. (2010) as shown in Figure S.1j,k |

As the number of parasitic salmon lice per fish is a model output that salmon farmers may observe, we calculated whether the number of parasites per fish should increase year round, and find four sites where this is the case: Chile (CH), Ireland (IM3 and IM4) and the site in the Broughton Archipelago (BCB) (Figure 3d). The remaining sites experience increasing number of parasites for only a percentage of the year: BCC (84%) BCV (76%), NLI (63%), NIN (57%), NB (54%), (NL) (45%), and NS (43%) (Figure 3d).

The follow-up treatment window varies seasonally and between regions (Figure 4a). The sites in Atlantic Canada (NB, NS and NL) and Lista, Norway (NLI) show large seasonal variation in the best timing of follow-up treatments as there exists no timing that will fall within the treatment window for all times of the year (Figure 4a, third row: no horizontal line can be drawn between the curves for *T*_1_ and *T*_2_). Temperature patterns determine the timing of the follow-up treatments, and the equations describing the beginning and end of the treatment windows (equations (3) and (4)), which assume that past temperatures are equal to the current temperature, are a reasonably good approximation when temperatures are not strongly seasonal (Figure 4b). At any given temperature, if temperatures have historically been warmer maturation occurs more quickly, and visa versa, and these historical effects in seasonal environments are the reason for the disparity between the approximation when past and current temperatures are assumed to be the same (Figure 4b, black dashed line), and the follow-up treatment window when past temperatures are allowed to reflect seasonality (Figure 4b, colors). Figure 4b shows that at warmer temperatures the treatment window narrows as chalimus maturation (equation (4)) occurs nearly as quickly as larval maturation and attachment (equation (3)). The maximum sea surface temperature observed in our dataset was 16°C and at this temperature the treatment window, assuming past and current temperatures are equal, is 12 to 14 days after the initial treatment.

## 4. Discussion

We have found that regional differences in climate can translate into substantial differences in population dynamics, which are important considerations for salmon lice management in different regions. Our analyses consider seasonal environments and we find that ignoring such seasonality can substantially bias prediction (Figure 2e,f). The dynamics of salmon lice in Chile and Ireland (CH, IM3, and IM4) are similar; these are the warmest sites with the fastest rates of population growth and the lowest stocking densities (Figure 2a,b). The sites in Atlantic Canada and Norway (NS, NB, NL, NIN and NLI) have the lowest temperatures, the slowest rates of population growth, and the highest stocking densities (Figure 2a,b), with the exception of Lista, Norway (NLI), which experiences a wide range of temperatures (Figure 1) and has more intermediate growth rates and critical stocking densities (Figure 2a,b). These sites with colder winter temperatures (NS, NB, NL, NIN and NLI) experience population declines during the winter, although the population rebounds in the summer and increases in size from year-to-year (Figure 2d).

Our model predictions agree with the critical stocking density of salmon for the Broughton Archipelago reported in Frazer et al. (2012) (Table 1). The models used in Frazer et al. (2012) ignore seasonality, but we found that the predicted critical stocking density for the Broughton Archipelago site (BCB) was relatively insensitive to the inclusion of seasonality in the model formulation (Figure 2f). The stage structure predicted by our model (equations (1) and (2)) tends to overestimate the abundance of immature stages relative to more mature stages, with the exception of nauplii relative to copepodids (Table 1), however, data on the stage structure of salmon lice populations are limited and Byrne et al. (2008) suggests that the relatively low density of larvae observed relative to the predictions of mathematical models may be due to the immediate dispersal of larvae following hatching.

The follow-up treatment windows that our analyses suggest are consistent with expert knowledge. In a modelling study utilizing the SLiDESim framework, Robbins et al. (2010) found that treatments occurring in pairs, with a follow-up treatment occurring six weeks after the initial treatment, is an effective approach to salmon lice management in Scotland, while anecdotally, it was thought that follow-up treatments after three to four weeks were best (Robbins et al., 2010). The nearest sites that we studied are IM3 and IM4 west of Ireland, and we found that follow-up treatments timed between 12 and 26 days will insure all larvae that escaped the initial treatment are in the parasite stage, but not yet having reached the reproductive adult female stage, when the follow-up treatment occurs. As such, our results are in approximate agreement with the anecdotal follow-up treatment timing reported for Scotland: a follow-up treatment after 3 weeks would always fall within the follow-up treatment window we recommend for IM3 and IM4 (Figure 4a). All of our results pertaining to treatment window recommendations are direct consequences of the *γ*_*P*_ (*T*) and *γ*_*C*_ (*T*) functions, which are parameterized from the available data on the maturation rates of nauplii and chalimi (Figure S.1e and as can be observed directly from equations (3) and (4)).

Despite the agreement of our model predictions with some observations, for the Newfoundland site with the empirically determined value of salinity, our model predicts that salmon lice should not be present (Figure 2a, *µ* = −0.02), which contradicts reality. We attribute this result to the observed low salinity (Figure 1g), because when higher values of salinity are used the model predicts that salmon lice will increase (Figure 2a, see (NL)). The low salinity values have been reported elsewhere (Rittenhouse et al., 2016) and are likely due to freshwater discharge from a hydroelectric generating station at the head of Bay D’Espoir (Brewer-Dalton et al., 2015). The most likely explanation for the failure of the model to predict salmon lice persistence when salinity is low, may be the failure of our model to account for short term (i.e., daily) fluctuations in salinity as previously highlighted in Groner et al. (2016).

Our analysis seeks to understand how different local temperature and salinity conditions will affect salmon lice population dynamics, and to facilitate this comparison we keep all other factors constant across regions. The abundance of host fish, species of host, and competition between salmon lice species varies regionally and will likely affect local dynamics (Costello, 2006). For any one site of interest, the estimated Floquet exponent, *µ*, or the critical stocking density, *f*_*crit*_, could be more accurately estimated by including some of these factors. In Section S.1 we discuss the parameterization of our life history functions given the available data from different salmon lice species and local adaptation, and we note that the dominant salmon louse species/subspecies varies between regions (Costello, 2006).

We measured population growth using two different quantities: the Floquet exponent, *µ*, and the basic reproductive ratio, *R*_0_. These two quantities measure, respectively, asymptotic year-to-year and asymptotic generation-to-generation change in population size. While the Floquet exponent has a clear applied relevance, *R*_0_ may be useful to measure the level of investment required to manage salmon lice. For example, for ordinary differential equation models the fraction of individuals that must be vaccinated to eradicate a disease is related to *R*_0_ (Scherer and McLean, 2002), but establishing such a relationship for more complex dynamical systems, and more general approaches to pest management, remains an open problem.

We have studied the regional differences in salmon lice population dynamics to better understand differences in regional management, however, future research is needed to address this challenge directly. We find that in Atlantic Canada and Norway during the winter months salmon lice abundance will decline, with the larval, chalimus, and pre-adult stages particularly affected. This decline in the abundance of all stages, as well as the bottlenecked stage structure, may imply different best timings of chemotherapeutic treatments in regions that experience cold winters, and our discussion of the regional differences in salmon lice population dynamics will be useful in interpreting the basis for these regionally specific management recommendations.

## Acknowledgements

The salinity data for Bay d’Espoir, NL was provided by Mathieu Ouellet (Marine Environmental Data Section, Oceans Science Branch, Department of Fisheries and Oceans Canada). AH would like to thank Martin Krkosek, Mark A. Lewis, Andry Ratsimandresy and James Watmough for helpful suggestions. This work was supported by the Atlantic Association for Research in the Mathematical Sciences and a National Sciences and Engineering Discovery Grant (RGPIN 2014-05413) to AH and an NSERC Grant to X.-Q. Zhao.

## Supplementary Online Material

This Supplementary Online Material (SOM) consists of five sections: S.1 Parameter descriptions and estimation; S.2 Model derivation; S.3 Threshold conditions for population growth; S.4 Numerical methods; and S.5 The follow-up treatment window.

### S.1 Parameter descriptions and estimation

Descriptions of all parameters are provided in Table S.1. The model (equations (1) and (2) in the main text) does not explicitly consider inflow and outflow of larval stages from the salmon netpens due to ocean currents, but we note that outflow likely exceeds inflow, such that the net effect is loss of larval lice from the salmon netpens. We assume that larvae that enter and leave the netpens have identical levels of development, such that larval net flow can be understood as being incorporated into two parameters: the mortality/loss rate of nauplii, *µ*_*P*_ (*t*), is considered to incorporate the net outflow of nauplii, such that if ocean currents are strong then *µ*_*P*_ (*t*) is larger, and the per fish attachment rate of copepodids, *ι*, is considered to incorporate the net outflow of copepodids, such that if ocean currents are strong, then *ι* is smaller. This approach to modelling larval flow is oversimplistic, but a more realistic formulation would require a substantial increase in model complexity, additional data, and it is not clear that this additional realism is necessary for our study.

Temperature and salinity are variable between sites, but the functions that describe the dependence of the model parameters on temperature, *T*, and salinity (measured in practical salinity units, psu), *S*, are the same across sites. The equations for life history parameters as a function of temperature and salinity were estimated by fitting curves to the laboratory results and the assumed functional forms were based on previous studies (Stien et al., 2005; Rittenhouse et al., 2016; Samsing et al., 2016).

Our model parameterization focuses on the *Lepeophtheirus salmonis* species of salmon lice. *L. salmonis* consists of two subspecies: *salmonis*, found in the Atlantic ocean, and *oncorhynchi*, found in the Pacific Ocean. These subspecies are genetically different, although, less so than different species (Skern-Mauritzen et al., 2014), and are capable of reproducing, but do not do so due to geographic barriers. It is not known whether these genetic differences at the subspecies level affect life history parameters (Elmoslemany et al., 2015), but a prior study (Stien et al., 2005) aggregates data from both subspecies. It is not possible to fully parameterize our model for one subspecies because not all relationships have been studied for both subspecies. Laboratory data that used the *salmonis* (black symbols) and *oncorhynchi* (blue symbols) subspecies are indicated in Figure S.1 and data from both subspecies of *L. salmonis* were combined to estimate the statistical fits.

Laboratory data for *Caligus rogercresseyi*, the salmon louse species that is dominant in Chile (Costello, 2006), are shown in green in Figure S.1. The *C. rogercresseyi* data was not considered for the statistical fits, but is shown for comparison. *C. rogercresseyi* do not have a pre-adult stage, but our modelling framework (equations (1) and (2)) aggregates both the chalimi and pre-adult stages into the attached, non-reproductive *C*(*t*)-stage, and so our model is an appropriate framework for *C. rogercresseyi*.

For all parameters related to egg hatching, *υ*(*T, S*), *η*(*T*), and *ϵ*(*T*), the data from Samsing et al. (2016 - crosses) may be most reliable since adult female lice were allowed to remain attached to their host salmonid during this study, whereas adult female salmon lice were removed from their host fish and transported to a laboratory for all other studies (Johnson and Albright, 1991; Boxaspen and Naess, 2000; Heutch et al., 2000; Bravo, 2010).

**Table S.1.** Description of model parameters.

| Parameters | Description | Dimensions |
| --- | --- | --- |
| $\iota$ | Attachment rate | $\text{time}^{-1}\text{number}^{-1}$ |
| $f$ | Number of fish per farm | number |
| $\eta(t)$ | Eggs per clutch | unitless |
| $\epsilon(t)$ | Egg string production rate | $\text{time}^{-1}$ |
| $v(t)$ | Proportion of eggs that produce viable nauplii | unitless |
| $\mu_P(t)$ | Mortality rate of nauplii | $\text{time}^{-1}$ |
| $\mu_I(t)$ | Net loss rate of copepodids | $\text{time}^{-1}$ |
| $\mu_C(t)$ | Mortality rate of chalimi and pre-adults | $\text{time}^{-1}$ |
| $\mu_A(t)$ | Mortality rate of adult females | $\text{time}^{-1}$ |
| $\gamma_P(t)$ | The maturation rate of nauplii | $\text{time}^{-1}$ |
| $\gamma_C(t)$ | The maturation rate of chalimi and pre-adults | $\text{time}^{-1}$ |
| $\tau_P(t)$ | The time taken to complete the nauplii stage for nauplii that mature to the copepodid stage at time, $t$ | time |
| $\tau_C(t)$ | The time taken to complete the chalimi and pre-adult stages for louse that mature to the adult female stage at time, $t$ | time |

The viability of salmon lice eggs depends on both temperature and salinity, and is,

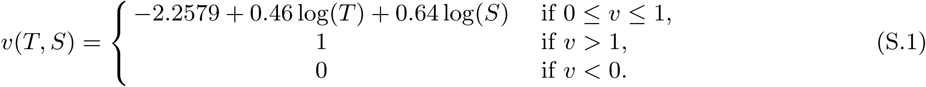

The data that describes the effect of temperature is from *L. salmonis salmonis* (Figure S.1a), while the data describing the effect of salinity is from *L. salmonis oncorhynchi* (Figure S.1b). Published data are insufficient to determine a temperature by salinity interaction since only fixed salinity is considered for a range of temperatures (Samsing et al., 2016), and fixed temperature is considered for a range of salinity (Johnson and Albright, 1991).

The number of eggs per string is,

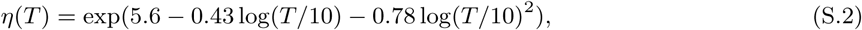

which is the fit described by Samsing et al. (2016) for *L. salmonis salmonis* (Figure S.1c). Data from Boxaspen and Naess (2000 - open squares) were excluded as they were not consistent with Samsing et al. (2016 - crosses). Bravo (2010) reports 31 eggs per string for *C. rogercresseyi* during an experiment where temperatures ranged from 10 - 18.5°C, and explains that this lower number of eggs per string may be because *C. rogercresseyi* is three times smaller than *L. salmonis*. Costello (2006) states that the number of eggs per string ranges from 100 to 1000 and depends on the time of year, host species, and louse species and size, also stating that 500 eggs per female louse is likely an underestimate for louse attached to farmed salmonids.

The rate of egg string production is,

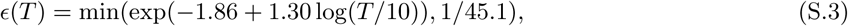

where the minimum is taken to prevent extrapolation beyond the minimum temperature observed experimentally. The fit is based on data which are a mix of the two *L. salmonis* subspecies, but these data from different subspecies agree quite closely (Figure S.1d, blue and black symbols). Data for *C. rogercresseyi* (Figure S.1d, green squares containing an ×, Montory et al. 2018) was not considered for the model fit, but shows that egg strings are produced more quickly for *C. rogercresseyi* at 5-10°C as compared to *L. salmonis*.

The maturation rate of nauplii to copepodids is,

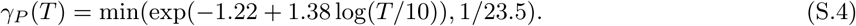

The minimum is taken to prevent extrapolation below 2°C, the lowest temperature for which nauplii maturation experiments were performed. The function *γ*_*P*_ (*T*) was also estimated in Stien et al. (2005), however, the fit described in equation (S.4) also considers more recent data collected by Samsing et al. (2016, Figure S.1e, crosses). While some data for the *oncorhynchi* subspecies (Figure S.1e, blue squares, Johnson and Albright 1991) are included in the model fit, these data are consistent with the data for the *salmonis* subspecies (Figure S.1e, black symbols). The maturation rate of nauplii for *C. rogercresseyi* (Figure S.1e, green squares with an *×*, Montory et al. 2018) is also consistent with *L. salmonis* (black symbols).

For the rate of maturation from chalimus to adult female, we use the fitted values of Stien et al. (2005),

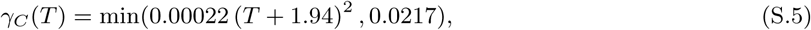

where the minimum is taken to prevent extrapolation below 8°C, the lowest temperature for which chalimi maturation rates were measured (Figure 4c in Stien et al. 2005; Figure S.1e in this SOM). The data measuring the maturation rate of chalimi for *C. rogercresseyi* (Figure S.1e, green circles with a +, Gonzalez and Carvajal 2003) are consistent with the fitted relationship from Figure 4c in Stien et al. 2005, which relies on data from laboratory experiments that used *L. salmonis salmonis*.

The mortality rates of nauplii, copepodid, chalimus and adults are assumed to be a function of salinity. For nauplii,

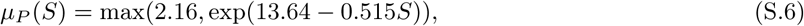

(see Figure S.1f). This function assumes that nauplii survive half a day at low salinities and becomes very large for high salinities consistent with the observation that all nauplii survived at 34 psu (Samsing et al., 2016). Samsing et al.’s observation is from *L. salmonis salmonis*, but all other observations are for the *oncorhynchi* subspecies.

For copepodids there is some evidence of a relationship between temperature and mortality in *C. rogercresseyi* (Figure S.1g, Montory et al. 2018). These data are shown to comprehensively report all data we assembled to parameterize our model, but our model assumes no temperature-mortality relationship because no studies for *L. salmonis* found this relationship.

For copepodids, the salinity-mortality relationship that we assume is,

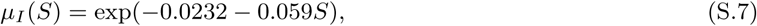

(see Figure S.1h). These data are a mix of the two *L. salmonis* subspecies. The data from Bricknell et al. (2006, open diamonds) was excluded from the fit because these data were not consistent with Tucker et al. (2002, solid circles) and Samsing et al. (2016, crosses) at 32 psu. For lower salinities, other than Bricknell et al. (2006, open diamonds) which was excluded, the only available data is from Johnson and Albright (1991, blue solid square) and these data are for the *oncorhynchi* subspecies. For *C. rogercresseyi*, data from Montory et al. (2018) shows that mortality rates are low (Figure S.1h, green square with an *×*) relative to *L. salmonis* at 34 psu.

Based on Connors et al. 2008 (Figure S.1h, blue inverted triangles), the relationship between salinity and the mortality rate for the parasitic salmon lice stages is,

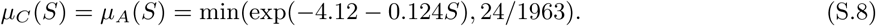

The experiments of Connors et al. (2008) did not distinguish between the mortality rates of chalimi and adult salmon lice and so we set *µ*_*C*_ (*S*) = *µ*_*A*_(*S*). In addition, the highest salinity considered by Connors et al. (2008) was 28 psu at which salmon lice survived 1963 hours: the salinity in salmon aquaculture pens at our sites may approach 35 psu, and to prevent extrapolation of the mortality function beyond what was observed we set a lower bound on the per capita mortality rate at 24/1963 lice per day. The estimates of *µ*_*A*_(*S*) for *C. rogercresseyi* (Figure S.1h, green squares containing a triangle) are from fitting an exponential decay function to the adult survival data in Table 1 of Bravo et al. (2008), however, these data are for adult sea lice removed from a host fish with subsequent survival monitored for 24 hours, while Connors et al.’s methodology considers parasitic *L. salmonis oncorhynchi* attached to pink *Oncorhynchus gorbuscha* and chum *O. keta* salmon monitored for > 1963 hours. Therefore, the Connors et al. (2008) data was considered more accurate. Nonetheless, *Caligus elogatus* are more sensitive to salinity than either *L. salmonis* or *C. rogercresseyi* as Landsberg et al. (1991) found that after 20 minutes at 8.2 psu none of the *C. elogatus* which had been attached to red drum (*Sciaenops ocellatus*) were still alive. In contrast, after 30 mins at 10 psu, 95-100% of *C. rogercresseyi* were still alive, with this percentage dropping to 15-45% after 24 hours (Bravo et al., 2008). Furthermore, the experiments of Connors et al. (2008) suggest that at 8.2 psu, 50% of adult *L. salmonis* will still be alive after 7 days.

Laboratory experiments have investigated the relationship between temperature and the copepodid attachment rate, *ι*, however no clear relationship was demonstrated (Figure S.1j). Bricknell et al. (2006) reports a relationship between salinity and the number of copepodids attached to salmon, however, the data describing natural mortality during this experiment are insufficient to allow us to estimate *ι*. We also expect that ocean currents may have a strong effect on copepodid attachment rates and these effects may be more important than temperature and salinity.

Only the BCB site contained sufficient data to estimate the number of fish on a farm, *f*, and the copepodid attachment rate, *ι*. Our mathematical model assumes a constant number of fish and no chemotherapeutic treatments and the BCB site satisfies these assumptions between March 3, 2003 to January 4, 2004. To estimate *f*, we calculated the mean number of fish on the BCB farm during the aforementioned period (506,737). During this same time period, we used data describing the number of *L. salmonis oncorhynchi* chalimi, pre-adults, and adult females, and performed maximum likelihood assuming a normal distribution of error to estimate the attachment rate with seasonality, *ι* = 2.1 *×* 10^−9^ (Figure S.1j,k, solid line), and ignoring seasonality, *ι* as 2.4 *×* 10^−9^ (Figure S.1j,k, dashed line), by considering only the average temperature (*a* = 8.3°C).

**Fig. S.1.**
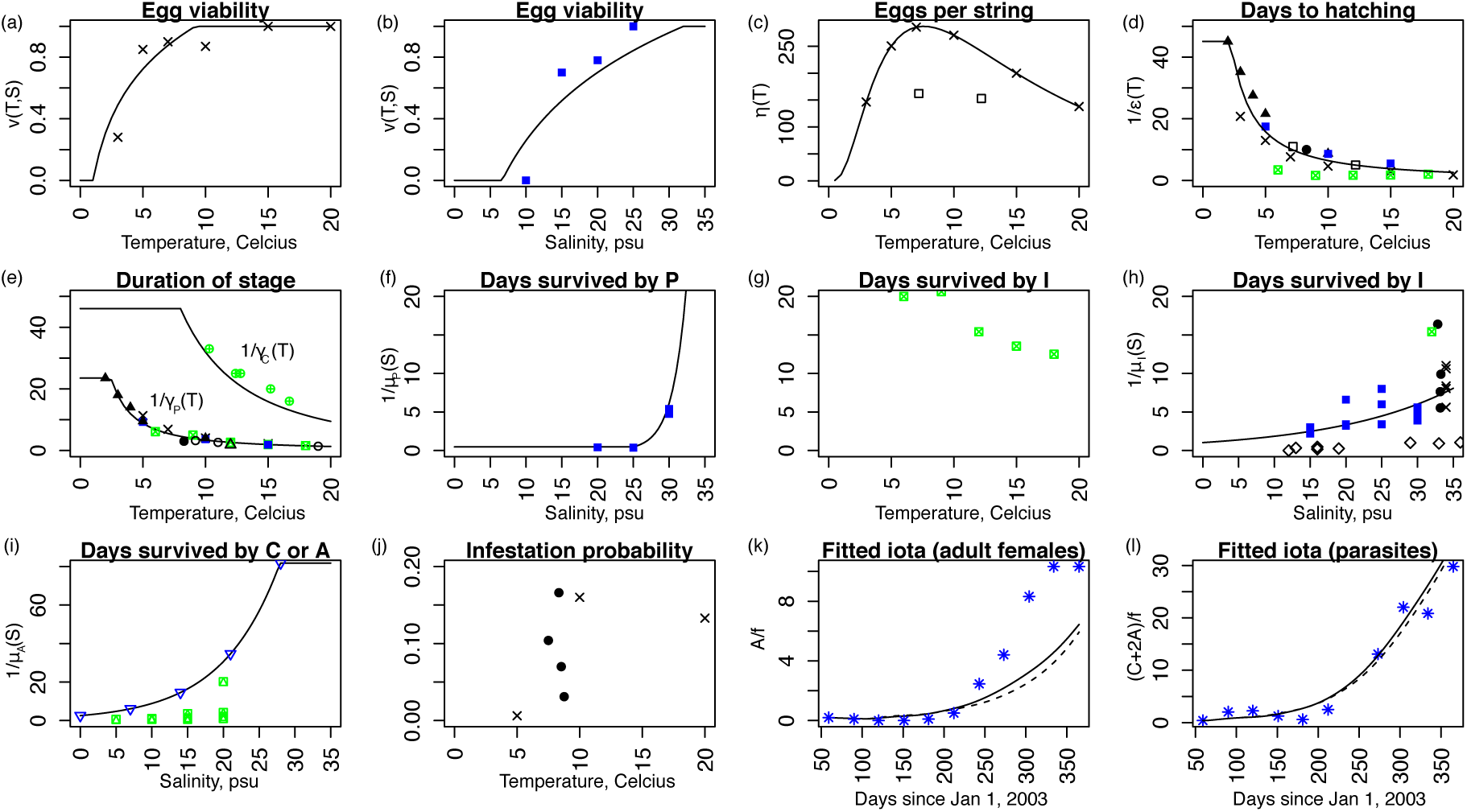
Published laboratory studies describe the relationship between life history parameters and temperature and salinity. We fitted functions (lines) to data from published laboratory studies (symbols) with *Lepeophtheirus salmonis salmonis* (black) and *L. salmonis oncorhynchi* (blue). Data from published laboratory studies with *Caligus rogercresseyi* are shown with green symbols: these data are for comparison and were not used for model fits. Black symbols (*L. salmonis salmonis*): Samsing et al. 2016 - crosses; Heutch et al. 2000 - open squares; Boxaspen and Naess 2000 - solid triangles; Tucker 2002 - solid circles; Johanessen 1978 - open circles; Wootten et al. 1982 - open triangles; and Bricknell et al. 2006 - open diamonds. Blue symbols (*L. salmonis oncorhynchi*): Johnson and Albright 1991 - solid squares; Connors et al. 2008 - open inverted triangles; and Marty et al. 2010 - astericks. Green symbols: Montory et al. 2018 - square with ×; Bravo et al. 2008 square with triangle; and Gonzalez and Carvajal 2003 - circle with +. Full details of the model parameterization and the equations are found in Section S.1.

#### S.1.1 Local adaptation

In Table S.2, we list all the data sources for the parameterization of each of the functions equations (S.1) - (S.8), as well as the origin of the salmon lice and the approximate year of the experiment. This is pertinent information to assess any bias that could arise due to local adaptation since our analyses apply these parameter estimates worldwide. We investigate local adaptation directly for the nauplius maturation rate, *γ*_*P*_ (*T*), because salmon lice of varied geographic origins contribute to this parameter estimate. In Figure S.2, we consider the source of origin of the *L. salmonis* in relation to local temperatures and the fitted *γ*_*P*_ (*T*) curve. We consider salmon lice ‘adapted’ if the temperature of the laboratory experiment (Figure S.2, the horizontal position of the symbols) is within the temperature range for the region of origin for the salmon lice (Figure S.2, the horizontal range of the bolded portion of the curve), and visa versa for ‘not adapted’. Firstly, we note that adapted and not adapted salmon lice have recorded near identical number of days in the nauplius stage prior to maturation, for example, in British Columbia salmon lice are not adapted at 15°C, but mature after the same number of days as lice form Norway which are adapted at this temperature. Secondly, the experiments are not performed at extreme temperates beyond the natural ranges that the salmon lice are adapted to: only lice originating from Norway have been exposed to very cold temperatures during experiments, and in our model, only populations from Norway (and Atlantic Canada where source populations would also be adapted to cold temperatures) are simulated at these low temperatures. Therefore, the data used to parameterize *γ*_*P*_ (*T*) do not suggest that local adaptation is biasing our results.

**Fig. S.2.**
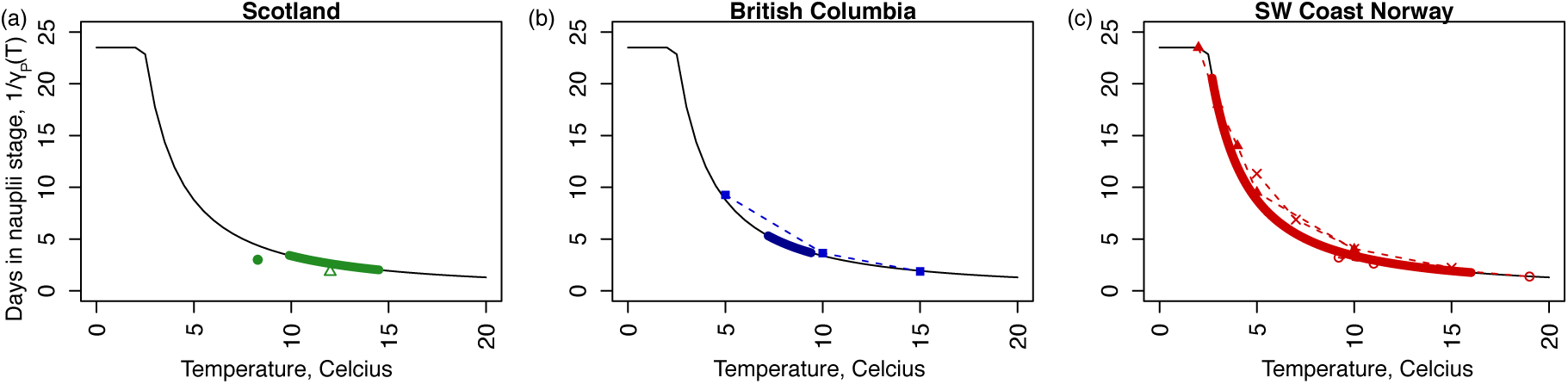
Local adaptation in *L. salmonis* does not invalidate our parameter estimates. We consider the origin of *L. salmonis* in relation to experimental data describing the number of days in the nauplii stage until maturation. We consider laboratory experiments where *L. salmonis* originate from a) Scotland (green symbols), b) the east coast of Vancouver Island, British Columbia (blue symbols), and c) Southwest Norway (green symbols). The temperatures for these experiments is compared to the range of sea surface temperatures from a nearby region (bold coloured portion of the curve), which corresponds to: a) Northern Ireland (IM4), b) Broughton Archipelago, British Columbia (BCB), and c) Lista, Norway (NLI). The fitted 1*/γ*_*P*_ (*T*) curve is shown in black. The experiments are performed at temperatures close to the local temperatures experienced by the salmon lice. Salmon lice exposed to temperatures outside the range they are adapted to record similar values to adapted salmon lice exposed to the same temperatures. The symbols are green: Tucker 2002 - solid circles and Wootten et al. 1982 - open triangles; blue: Johnson and Albright 1991 - solid squares; and red: Samsing et al. 2016 - crosses; Johanessen 1978 - open circles and Boxaspen and Naess 2000 - solid triangles.

**Table S.2.** The source of all data used to parameterize functions of temperature and salinity. All data are for *Lepeophtheirus salmonis* for the subspecies *salmonis* (no underline) and *oncorhynchi* (underline). Definitions of the parameter symbols are provided in Table S.1. Origin describes the place of origin for the salmon lice used for the experiments. The year is the publication year, unless the year of the experiment is specified (denoted with *). Range is the range of temperature or salinity values over which measurements are taken. For Boxaspen and Naess (2000), the origin of the salmon lice was not stated, but the text suggested the origin was Norway (denoted with **).

| Parameter | Origin | Year | Range (°C or psu) | Reference |
| --- | --- | --- | --- | --- |
| $v(T)$ | SW coast Norway | 2015* | [3,20] | Samsing et al. (2016) |
| $\eta(T)$ | SW coast Norway | 2015* | [3,20] | Samsing et al. (2016) |
| $\epsilon(T)$ | <u>E coast Vancouver Island, Canada</u> | 1991 | [5,15] | Johnson and Albright (1991) |
|  | SW coast Norway | 2015* | [3,20] | Samsing et al. (2016) |
|  | Norway? | 2000 | [2,10] | Boxaspen and Naess (2000) |
|  | W coast Norway | 2002 | [7.2,12.2] | Heutch et al. (2000) |
|  | W coast Scotland | 2002 | 8.3 | Tucker (2002) |
| $\gamma_P(T)$ | <u>E coast Vancouver Island, Canada</u> | 1991 | [5,15] | Johnson and Albright (1991) |
|  | SW coast Norway | 2015* | [5,15] | Samsing et al. (2016) |
|  | Norway** | 2000 | [2,10] | Boxaspen and Naess (2000) |
|  | W coast Scotland | 2002 | 8.3 | Tucker (2002) |
|  | Norway/sea water aquaria at 9 and 11°C | 1978 | [9.2,19] | Johanessen (1978) |
|  | Scotland? | 1982 | 12°C | Wootten et al. (1982) |
| $\gamma_C(T)$ | W coast Norway | 1996 | [9,10] | Grimnes and Jakobsen (1996) |
|  | N Norway | 1997* | 9.7 | Bjorn et al. (1998) |
|  | Reared at 7.5°C/W coast Norway? | 2000 | 7-10 variable | Finstad et al. (2001) |
|  | W coast Scotland | 2002 | 8.3 | Tucker (2002) |
| $v(S)$ | <u>E coast Vancouver Island, Canada</u> | 1991 | [10,25] | Johnson and Albright (1991) |
| $\mu_P(S)$ | <u>E coast Vancouver Island, Canada</u> | 1991 | [20,30] | Johnson and Albright (1991) |
|  | SW coast Norway | 2015* | 34 | Samsing et al. (2016) |
| $\mu_I(S)$ | <u>E coast Vancouver Island, Canada</u> | 1991 | [15,30] | Johnson and Albright (1991) |
|  | SW coast Norway | 2015* | 34 | Samsing et al. (2016) |
|  | W coast Scotland | 2002 | [32.8,33.3] | Tucker (2002) |
| $\mu_C(S)$ | <u>Broughton Archipelago, Canada</u> | 2006* | [0,28] | Connors et al. (2008) |

#### S.1.2 Temperature and salinity

We assume that temperature and salinity are seasonal and described by,

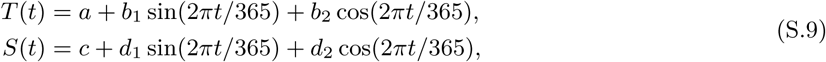

where *a* is the mean sea surface temperature in degrees Celcius (°*C*), *c*, is the mean salinity in psu, *b*_1_ and *b*_2_ affect the magnitude and timing of annual temperature changes, and *d*_1_ and *d*_2_ affect the magnitude and timing of annual salinity changes. For the BCB and NL sites, the salinity data was not seasonal since fitting the *d*_1_ and *d*_2_ coefficients did not improve the proportion of the variance explained (*R*^2^) by more than 0.02, and so *d*_1_ and *d*_2_ were set to zero at these sites. For all the sites in Canada, both temperature and salinity data were available, but for the sites in Chile, Ireland, and Norway only temperature data was available; here we assume a constant salinity of 31 psu, which was the mean salinity for all sites with available data, but excluding the NL site where salinity was very low. To facilitate comparisons with the Northern Hemisphere sites, the temperate data from Region X, Chile was shifted by 182.5 days (Figure 1).

Temperature and salinity data as shown in Figure 1 were compiled from several sources as described below.

##### CH: Finfish farm near Puerto Montt, Region X, Chile

Data consists of sea surface temperatures recorded from June 2000 to February 2001 (Bravo, 2003). This was the only southern hemisphere site and data was shifted by 182.5 days to enable comparisons with other sites. No salinity was available for this region, so our analyses assume a constant salinity of 31 psu, which is the average salinity reported for the BCB, BCC, BCV, NS and NB sites.

##### IM3: Weather buoy west of Ireland

Mean monthly temperature recorded from 2003-2013 at the M3 weather buoy west of Ireland (Dabrowski et al., 2016). No salinity data was available for this region, so our analyses assume a constant salinity of 31 psu.

##### IM4: Weather buoy west of Ireland

Mean monthly temperature recorded from 2003-2013 at the M4 weather buoy west of Ireland (Dabrowski et al., 2016). No salinity was available for this region, so our analyses assume a constant salinity of 31 psu.

##### BCB: Finfish farm in the Broughton Archipelago, British Columbia, Canada

Temperature and salinity data is from farm 24 in Marty et al. (2010) collected Jan 1, 2001 - Nov 1, 2007. Salinity at this site is consistent with an ‘oceanic’ site as described by Groner et al. (2016).

##### BCC: Lighthouse on the Central Coast of British Columbia, Canada

Mean monthly temperature and salinity (1954-2011) recorded at the McInnes Island lighthouse. Data is from Figure 3 in Brewer-Dalton et al. (2015).

##### BCV: Lighthouses on the west coast of Vancouver Island, BC

Mean monthly temperature and salinity (1935-2012) recorded at the Amphitrite Point and Kains Island lighthouses. Data is from Figure 2 in Brewer-Dalton et al. (2015).

##### NB: Hydrographic station in the mouth of the Bay of Fundy, NB

Mean monthly temperature and salinity (1971-2000) recorded at the Prince 5 hydrographic station. Data is from Figures 25 and 28 in Brewer-Dalton et al. (2015).

##### NS: Hydrographic station located off Halifax, Nova Scotia, Canada

Mean monthly temperature and salinity (1971-2000) recorded at the Station 2 hydrographic station located off Halifax, Nova Scotia. Data is from Figures 25 and 28 in Brewer-Dalton et al. (2015).

##### NL: Hermitage Bay-Bay d’Espoir, Newfoundland, Canada

Mean monthly temperature is from Figure 8 of Department of Fisheries and Oceans (2016). Monthly salinity was provided by the Department of Fisheries and Oceans (see Acknowledgements), but similar data is shown in Figure 18 in Brewer-Dalton et al. (2015). Salinity data with no associated depth was removed and only measurements taken between for 0-5m were included in the analysis. These salinity data were collected between 1994 and 2009.

##### NIN: Meterological station in Ingøy, Norway

Mean temperature measured every 14 days at a meterological station since 1942. Data is from Figure 1 in Samsing et al. (2016). No salinity data was available for this region, so our analyses assume a constant salinity of 31 psu.

##### NLI: Meterological station in Lista, Norway

Mean temperature measured every 14 days at a meterological station since 1942. Data is from Figure 1 in Samsing et al. (2016). No salinity data was available for this region, so our analyses assume a constant salinity of 31 psu.

**Table S.3.**
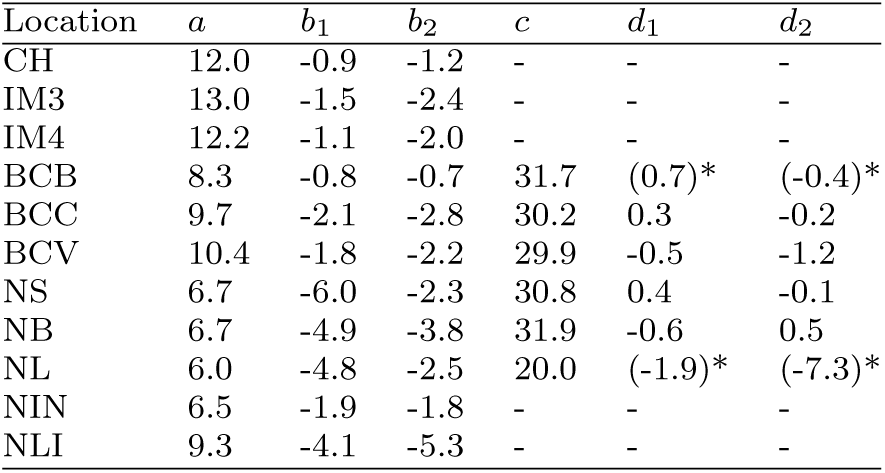
Site specific temperature and salinity parameter estimates. Parameters are described in equations (S.9): notably, *a* is the mean temperature and *c* is the mean salinity. Where the inclusion of *d*_1_ and *d*_2_ improves *R*^2^ by less than 0.02, this is indicated by * and parentheses indicate that these values were set to zero for the analyses. Where parameters cannot be calculated due to unavailable data this is indicated by -.

### S.2 Model derivation

Here we derive the mathematical model describing salmon lice dynamics (equations (1) and (2)). This derivation is provided in Rittenhouse et al. (2016), but in this section we provide some additional details, which will be useful for some readers. One minor difference between our model and that of Rittenhouse et al. (2016) is that our model has the number of eggs per string, *η*(*t*), and the egg string production rate, *ϵ*(*t*) depend on time.

Our mathematical model for salmon lice population dynamics incorporates a temperature-dependent maturation delay and salinity-dependent mortality rates. We note that temperature and salinity are time-dependent. Let *M*_*P*_ (*t*) be the maturation rate of nauplii, and *M*_*C*_ (*t*) be the maturation rate of chalimi/pre-adults. Assuming a 1:1 sex ratio, 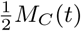 is the maturation rate of chalimi/pre-adults to the adult female stage. Then,

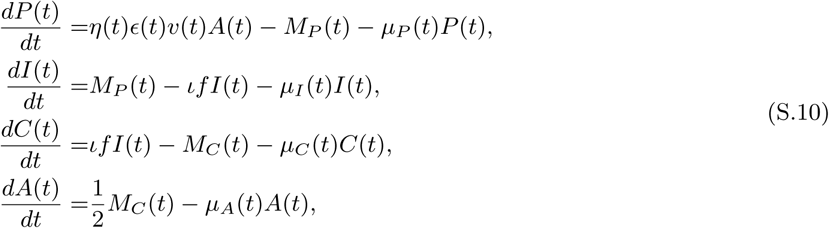

where the variables *P* (*t*), *I*(*t*), *C*(*t*) and *A*(*t*) are each salmon lice stage. All parameters are defined in Table S.1. To derive a model with time-delayed maturation after a threshold level of development is completed, we follow the model derived in Nisbet and Gurney (1983). Let *q* be the development level of salmon lice such that *q* increases at a temperature-dependent rate *γ*_*x*_(*T*(*t*)) = *γ*_*x*_(*t*) where *x* = *P* or *C*. Suppose *q* = *q*_*P*_ = 0 at the start of stage *P*, *q* = *q*_*I*_ at the transition from *P* to *I, q* = *q*_*C*_ at the transition from *I* to *C*, and *q* = *q*_*A*_ at the transition from *C* to *A*. Let *ρ*(*q, t*) be the density of salmon lice with development level *q* at time *t*. Then *M*_*P*_ (*t*) = *γ*_*P*_ (*t*)*ρ*(*q*_*I*_, *t*), *M*_*C*_ (*t*) = *γ*_*C*_ (*t*)*ρ*(*q*_*A*_, *t*).

Let *J*(*q, t*) be the flux, in the direction of increasing *q*, of salmon lice with development level *q* at time *t*. Then we have the equations (see, e.g., Kot 2001),

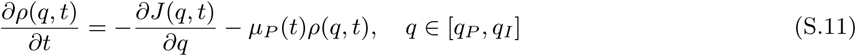

and,

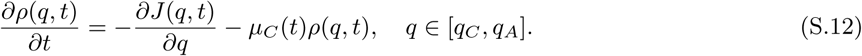

Since *J*(*q, t*) = *ρ*(*q, t*)*γ*_*x*_(*t*), with *x* = *P, C*, we have

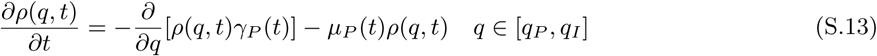

and,

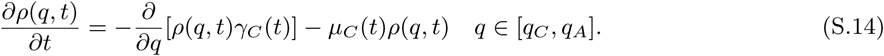

For the *P* state, system (S.13) has the boundary condition

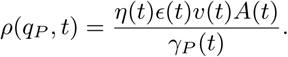

To solve system (S.13) with this boundary condition, we introduce a new variable

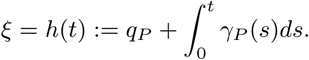

Let *h*^−1^(*ξ*) be the inverse function of *h*(*t*), and define

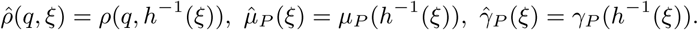

Given (S.13), we then have

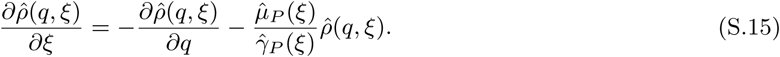

This equation is identical in form to the standard von Foerster equation (see Nisbet and Gurney 1982). Let 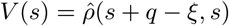. It follows from (S.15) that

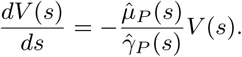

Since *ξ* − (*q* − *q*_*P*_) *≤ ξ*, we have

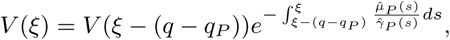

and hence,

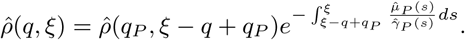

Define *τ*_*P*_ (*q, t*) to be the time taken to grow from development level *q*_*P*_ to level *q* by salmon lice who arrive at development level *q* at time *t*. Since 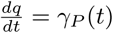 for *q* ∈ [*q*_*I*_, *q*_*P*_], it follows that

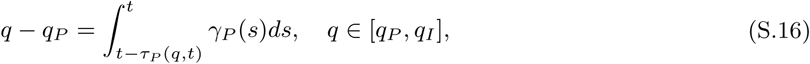

and hence,

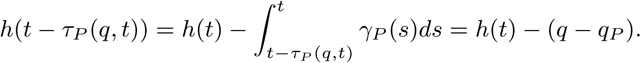

By the change of variable *s* = *h*(*α*), we then see that

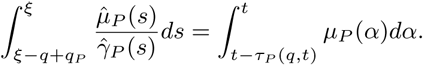

It follows that

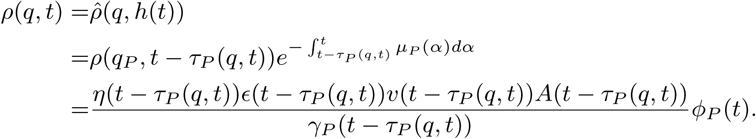

where 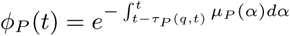. Denote *τ*_*P*_ (*t*) = *τ*_*P*_ (*q*_*I*_, *t*), then we have

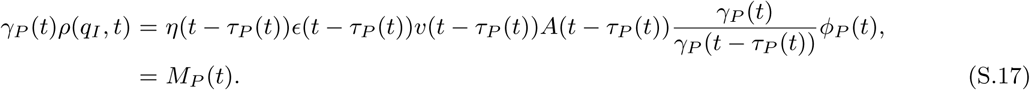

Substituting equation (S.17) into (S.10), we then have the 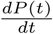 and 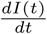 equations in the system (1) in the main text. By similar arguments, we can obtain,

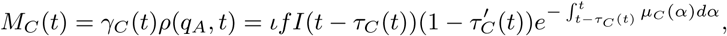

which when substituted into (S.10) gives the 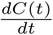 and 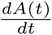 equations in system (1) in the main text.

We note that the maturation rates into the *I*(*t*) and *A*(*t*) stages (*M*_*P*_ (*t*) and *M*_*C*_ (*t*)), do not depend on the abundance of the preceding stages (*P* (*t*) and *C*(*t*)). For delay differential equation models, it is not always necessary to specify the dynamics of all stages, for example see Gurney et al. (1980), whereby the model (equation 6 in Gurney et al. 1980) describes the change in only the mature adult fly population. The parameter *T*_*D*_ is the time to reach the mature adult stage from an egg, such that the stage occurring between the egg and the mature adult stage (pupae) has been omitted from the model specification due to the assumption of time delayed maturation. For our model (equations 1), we include the *dP/dt* and *dC/dt* equations because we report results related to these quantities, however, the *dP/dt* and *dC/dt* equations do not affect the model dynamics.

Letting *q* = *q*_*I*_ in (S.16) we get,

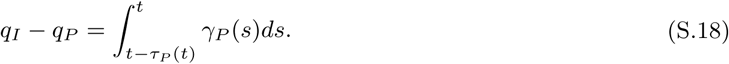

Taking the derivative with respect to *t* on both sides of (S.18) we obtain,

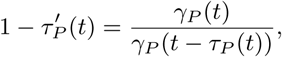

which is the 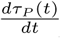 equation appearing in (2) in the main text. Define *τ*_*C*_ (*t*) to be the time taken to grow from development level *q*_*C*_ to level *q*_*A*_ by salmon lice who arrive at development level *q*_*A*_ at time *t*. We then have,

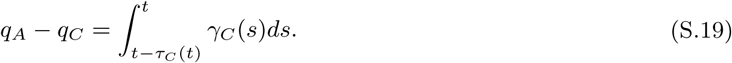

Taking the derivative with respect to *t* on both sides of (S.19) we have

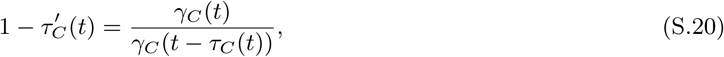

which is the 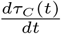 equation appearing in (2) in the main text. By virtue of (S.18) and (S.19), it easily follows that if *γ*_*x*_(*t*) is a periodic function, then so is *τ*_*x*_(*t*) with the same period (*x* = *P* or *C*).

### S.3 Threshold dynamics

In this section, we study the global dynamics of system (1) and (2). First, we will use the theory recently developed in Zhao (2017) to derive the basic reproduction ratio *R*_0_. Since the 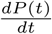 and 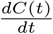 equations of system (1) are decoupled from the other equations, it suffices to study the following system:

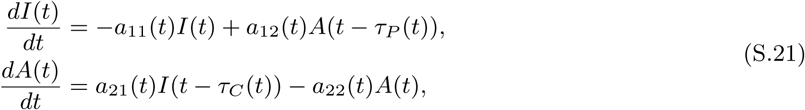

where 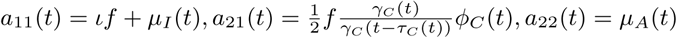, and 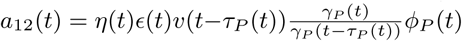. 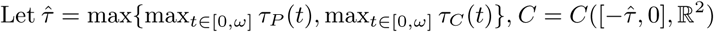. Then (*C, C*^+^) is an ordered Banach space equipped with the maximum norm and the positive cone *C*^+^. For any given continuous function 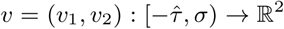 with *σ* > 0, we define *υ*_*t*_ ∈ *C* by

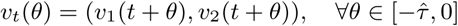

for any *t* ∈ [0, *σ*). Let *F* : ℝ → ℒ(*C*, ℝ^2^) be a map and *υ* (*t*) be a continuous 2 × 2 matrix function on ℝ defined as follows:

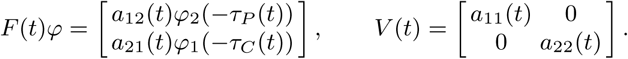

Then the internal evolution of the compartments *I* and *A* can be expressed by

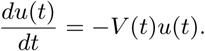

Let *Φ*(*t, s*), *t ≥ s*, be the evolution matrix of the above linear system. That is, *Φ*(*t, s*) satisfies

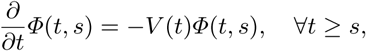

and

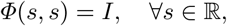

where *I* is the 2 × 2 identity matrix. It then easily follows that

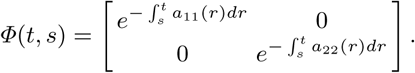

Let *C*_*ω*_ be the ordered Banach space of all continuous and *ω*-periodic functions from ℝ to ℝ^2^, which is equipped with the maximum norm and the positive cone 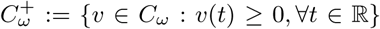. Suppose that *υ* ∈ *C*_*ω*_ is the initial distribution of copepodid and adult female individuals. Then for any given *s ≥* 0, *F* (*t* − *s*)*υ*_*t*−*s*_ is the distribution of offspring (copepodids and adult females) at time *t*− *s*, which is produced by the individuals who were introduced over the time interval 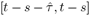. Then *Φ*(*t, t* − *s*)*F* (*t* − *s*)*υ*_*t*−*s*_ is the distribution of those individuals who newly entered the copepodid or adult female compartments at time *t* − *s* and remain in the compartments at time *t*. It follows that

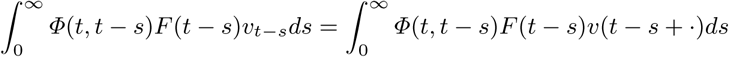

is the cumulative distribution of new copepodids and adult females at time *t* produced by all the copepodids and adult females introduced at all times prior to *t*.

Define a linear operator *L* : *C*_*ω*_ → *C*_*ω*_ by

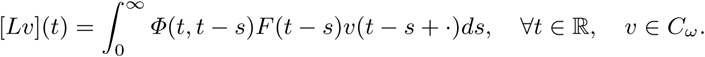

Following Zhao (2017), we define *R*_0_ = *r*(*L*), the spectral radius of *L*. Let 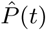 be the solution maps of system (S.21) on *C*, that is, 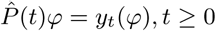, where *y*(*t, φ*) is the unique solution of (S.21) with *y*_0_ = *φ* ∈ *C*. Then 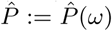 is the Poincaré map associated with linear system (S.21). Let 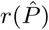 be the spectral radius of 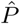. In light of Zhao (2017, Theorem 2.1), we have the following observation.

#### Lemma 1

*R*_0_ − 1 *has the same sign as* 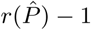.

By Hale (1993, Theorem 6.1.1) and Smith (1995, Theorem 5.2.1), we obtain the following result for linear system (S.21).

#### Lemma 2

*For any φ* ∈ *C*^+^, *system* (S.21) *has a unique solution y*(*t, φ*) *with y*_0_ = *φ, and y*(*t, φ*) ≥ 0 *for all t* ≥ 0.

Let

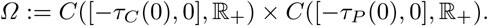

Then we further have the following result.

#### Lemma 3

*For any φ* ∈ *Ω, system* (S.21) *has a unique solution z*(*t, φ*) *with z*_0_ = *φ, and z*_*t*_(*φ*) := (*z*_1*t*_(*φ*), *z*_2*t*_(*φ*)) ∈ *Ω for all t* ≥ 0.

*Proof* 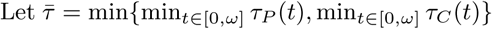. For any 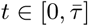, since *t*− *τ*_*P*_ (*t*) and *t* − *τ*_*C*_ (*t*) are strictly increasing in *t*, we have

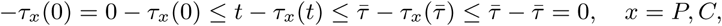

and hence,

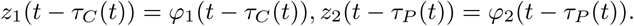

Therefore, we have the following ordinary differential equations for 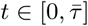:

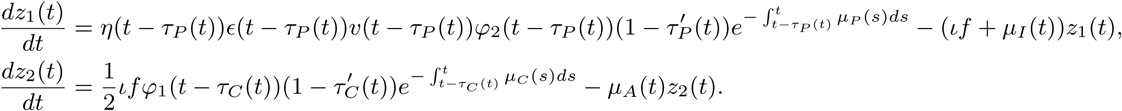

Given *φ* ∈ *Ω*, the solution (*z*_1_(*t*), *z*_2_(*t*)) of the above system exists for 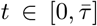. In other words, we have obtained values of *ψ*_1_(*θ*) = *z*_1_(*θ*) for 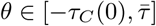 and *ψ*_2_(*θ*) = *z*_2_(*θ*) for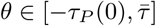.

For any 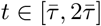, we have

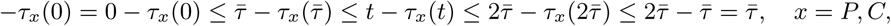

and hence, *z*_1_(*t-τ*_*C*_ (*t*)) = *ψ*_1_(*t-τ*_*C*_ (*t*)), *z*_2_(*t-τ*_*P*_ (*t*)) = *ψ*_2_(*t-τ*_*P*_ (*t*)). Solving the following system of ordinary differential equations for 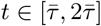 with 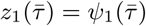 and 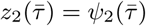:

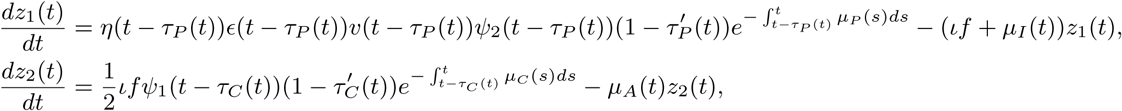

we then get the solution (*z*_1_(*t*), *z*_2_(*t*)) on 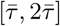. Repeating this procedure for 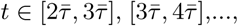 it then follows that for any *φ* ∈ *Ω*, system (S.21) has a unique solution *z*(*t, φ*) with *z*_0_ = *φ* and *z*_*t*_(*φ*) = (*z*_1*t*_(*φ*), *z*_2*t*_(*φ*)) ∈ *Ω* for all *t* ≥ 0.

#### Remark 1

By the uniqueness of solutions in Lemmas 2 and 3, it follows that for any *ψ* ∈ *C*_+_ and *ϕ* ∈ *Ω* with *ψ*_1_(*θ*) = *ϕ*_1_(*θ*) for all *θ* ∈ [*-τ*_*C*_ (0), 0] and *ψ*_2_(*θ*) = *ϕ*_2_(*θ*) for all *θ* ∈ [*-τ*_*P*_ (0), 0], we have *y*(*t, ψ*) = *z*(*t, ϕ*), *∀t* ≥ 0, where *y*(*t, ψ*) and *z*(*t, ϕ*) are solutions of system (S.21) satisfying *y*_0_ = *ψ* and *z*_0_ = *ϕ*, respectively.

Let *P* (*t*) be the solution maps of system (S.21) on *Ω*, that is, *P* (*t*)*φ* = *z*_*t*_(*φ*), *t* ≥ 0, where *z*(*t, φ*) is the unique solution of system (S.21) with *z*_0_ = *φ* ∈ *Ω*. By the arguments similar to those in Lou and Zhao (2017, Lemma 3.5), we have the following result.

#### Lemma 4

*P* (*t*) : *Ω* → *Ω is an ω-periodic semiflow in the sense that* (*i*) *P* (0) = *I;* (*ii*) *P* (*t* + *ω*) = *P* (*t*) ° *P* (*ω*), *∀t* ≥ 0; (*iii*) *P* (*t*)*φ is continuous in* (*t, φ*) ∈ [0, *∞*) *× Ω.*

Let *P* be the Poincaré map of the linear system (S.21) on the space *Ω*, and *r*(*P)* be its spectral radius. Then we have the following threshold result for system (S.21).

#### Lemma 5

*The following statements are valid:*

i. *If r*(*P) ≤* 1, *then* lim_*t*→*∞*_(*I*(*t, φ*), *A*(*t, φ*)) = (0, 0) *for any φ* ∈ *Ω.*
ii. *If r*(*P)* > 1, *then* lim_*t*→*∞*_(*I*(*t, φ*), *A*(*t, φ*)) = (*∞, ∞*) *for any φ* ∈ *Ω \ {*0*}.*

*Proof* For any given *φ, ψ* ∈ *Ω* with *φ* ≥ *ψ*, let ū (*t*) = *u*(*t, φ*) and *u*(*t*) = *u*(*t, ψ*) be the unique solutions of system (S.21) with *u*_0_ = *φ* and *u*_0_ = *ψ*, respectively. Let 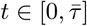.

Since for any 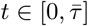,

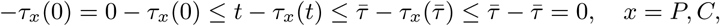

we have *ū*_1_(*t* − *τ*_*C*_ (*t*)) = *φ*_1_(*t* − *τ*_*C*_ (*t*)), *u*_1_(*t* − *τ*_*C*_ (*t*)) = *ψ*_1_(*t* − *τ*_*C*_ (*t*)), *ū*_2_(*t* − *τ*_*P*_ (*t*)) = *φ*_2_(*t* − *τ*_*P*_ (*t*)) and *u*_2_(*t − τ*_*P*_ (*t*)) = *ψ*_2_(*t-τ*_*P*_ (*t*)) for all 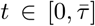, and hence, *ū*_1_(*t − τ*_*C*_ (*t*)) ≥ *u*_1_(*t − τ*_*C*_ (*t*)) and *ū*_2_(*t − τ*_*P*_ (*t*)) ≥ *u*_2_(*t − τ*_*P*_ (*t*)) for all 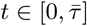. In view of *ū*(0) = *φ*(0) ≥ *ψ*(0) = *u*(0), the comparison theorem for cooperative ordinary differential systems implies that *ū*(*t*) ≥ *u*(*t*) for all 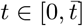. Repeating this procedure for 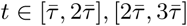, …, it follows that *u*(*t, φ*) ≥ *u*(*t, ψ*) for all *t* ∈ [0, *∞*). This implies that *P* (*t*) : *Ω* → *Ω* is monotone for each *t* ≥ 0. Next we show that the solution map *P* (*t*) : *Ω* → *Ω* is eventually strongly monotone. Let *φ, ψ* ∈ *Ω* satisfy *φ > ψ*. Denote 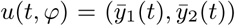 and *u*(*t, ψ*) = (*y*_1_(*t*), *y*_2_(*t*)). Without loss of generality, we assume that *φ*_1_ > *ψ*_1_.

Since 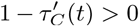, there exists a unique solution to the equation *t* − *τ*_*C*_ (*t*) = 0. Denote the unique solution of *t* − *τ*_*C*_ (*t*) = 0 as 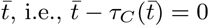.

*Claim 1.* There exists 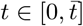 such that 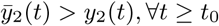.

We first prove that 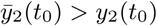 for some 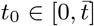. Otherwise, we have 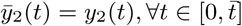, and hence, 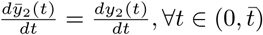. Thus, we have

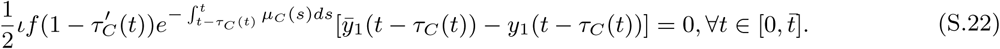

It follows that 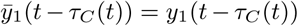 for all 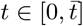, that is, *φ*_1_(*θ*) = *ψ*_1_(*θ*) for all *θ* ∈ [*-τ*_*C*_ (0), 0], which contradicts the assumption that *φ*_1_ > *ψ*_1_.

Let

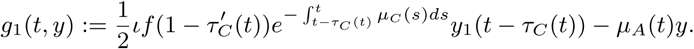

Since

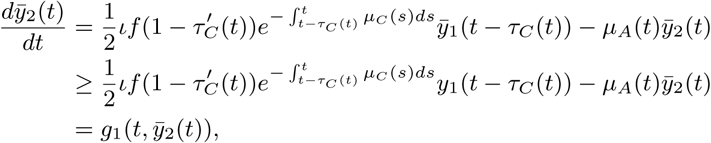

we have

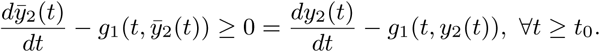

Since 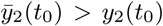, the comparison theorem for ordinary differential equations (Theorem 4, Walter 1997) implies that 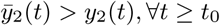.

Denote the unique solution to 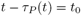 as 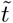.

*Claim 2.* 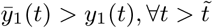.

Let

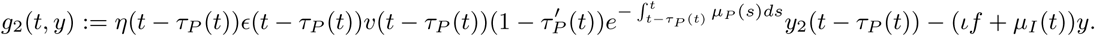

Then we have

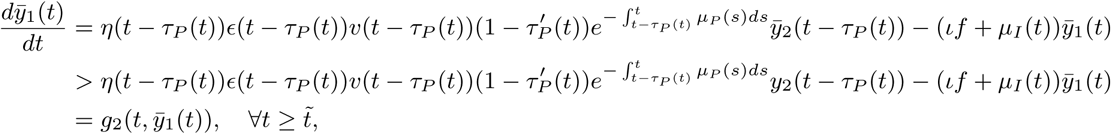

and hence,

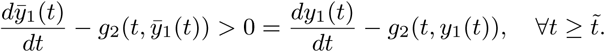

Since 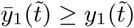, it follows from Walter (1997, Theorem 4) that 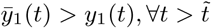.

In view of Claims 1 and 2, we obtain

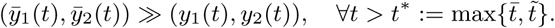

It follows that

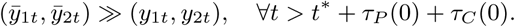

This shows that *P* (*t*) : *Ω* → *Ω* is strongly monotone for any *t* > *t*\* + *τ*_*P*_ (0) + *τ*_*C*_ (0). It follows from Hale (1993, Theorem 3.6.1) that the linear operator *P* (*t*) is compact on *Ω*. Choose an integer *n*_0_ such that *n*_0_*ω* > *t*\* + *τ*_*P*_ (0) + *τ*_*C*_ (0). Since 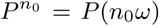, Liang and Zhao (2007, Lemma 3.1) implies that *r*(*P*) is a simple eigenvalue of *P* having a strongly positive eigenvector, and the modulus of any other eigenvalue is less than *r*(*P*). It then follows from Wang and Zhao (2017, Lemma 1) that there is a positive *ω*-periodic function 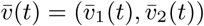 such that 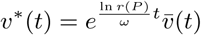 is a positive solution of system (S.21).

In the case where *r*(*P)* < 1, we have lim_*t*→∞_ *υ*\*(*t*) = 0. For any *φ* ∈ *Ω*, choose a sufficiently large number *K* > 0 such that 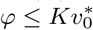. Then by the comparison theorem, we have

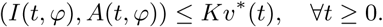

Hence, lim_*t*→∞_ *I*(*t, φ*) = lim_*t*→∞_ *A*(*t, φ*) = 0. This proves statement (i).

In the case where *r*(*P)* > 1, we have lim_*t*→∞_ *υ*\*(*t*) = ∞. For any *φ* ∈ *Ω* \ {0}, we have *u*_*t*_(*φ*) ≫ 0 for all *t* > *t* + *τ*_*P*_ (0) + *τ*_*C*_ (0). Without loss of generality, we assume that *φ* 0. Then we can choose a sufficiently small real number *δ* > 0 such that 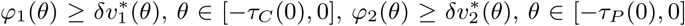. Then by the comparison theorem, we have

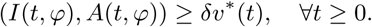

Hence, lim_*t*→∞_(*I*(*t, φ*), *A*(*t, φ*)) = (∞, ∞). This proves statement (ii).

By the same arguments as in Lou and Zhao (2017, Lemma 3.8), we have 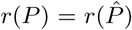. Combining Lemmas 1 and 5, we have the following result on the global dynamics of system (S.21).

#### Theorem 2

*The following statements are valid:*

i. *If R*_0_ < 1, *then the extinction equilibrium* (0, 0) *is globally attractive for system* (S.21) *in Ω;*
ii. *If R*_0_ > 1, *then all nontrivial solutions of system* (S.21) *go to infinity eventually.*

In the rest of this section, we derive the dynamics for the variables *P* (*t*) and *C*(*t*) in system (1). Under the compatibility condition

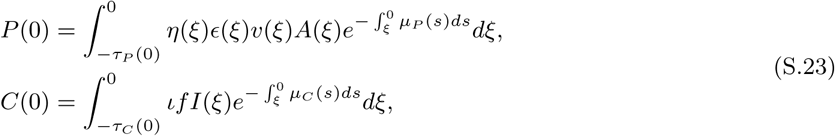

we can solve *P* (*t*) and *C*(*t*) as

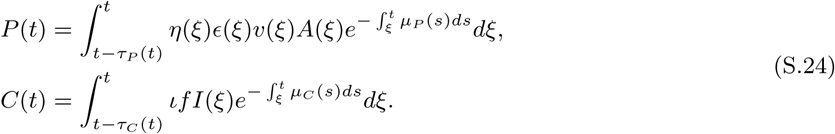

In the case where *R*_0_ < 1, we have lim_*t*→∞_ *A*(*t*) = lim_*t*→∞_ *I*(*t*) = 0, and hence, the expression (S.24) implies that lim_*t*→∞_ *P* (*t*) = lim_*t*→∞_ *C*(*t*) = 0.

In the case where *R*_0_ > 1, we obtain lim_*t*→∞_ *A*(*t*) = lim_*t*→∞_ *I*(*t*) = +∞. It then follows from (S.24) that lim_*t*→∞_ *P* (*t*) = lim_*t*→∞_ *C*(*t*) = +∞. Consequently, using these arguments combined with Lemma 1, we have the result stated in the main text (Theorem 1) describing the global dynamics of system (1) and (2).

### S.4 Numerical methods

The system of equations (1) and (2), was solved using the `dde()` and `pastvalue()` functions in the *PBSddesolve* package for R (Couture-Beil et al., 2016). This solver for delay differential equations is based on Simon Wood’s *solv95* program (Wood, 1999) written in C/C+ and using an adaptively stepped embedded RK2(3) scheme with cubic hermite interpolation of the lagged variables.

Figure 3 in the main text assumes *A*(0) = *f* and *P* (0) = *I*(0) = *C*(0) = 0 and Figure 4 assumes *A*(0) = *P* (0) = *I*(0) = *C*(0) = *f*. For Figures 3 and 4, we assumed the initial history, *t* < 0, for each variable was equal to their respective values at *t* = 0. The initial values of the time delays where calculated by numerically solving equation (S.18) with *q*_*I*_ − *q*_*P*_ = 1 and equation (S.19) with *q*_*A*_ − *q*_*C*_ = 1. To numerically evaluate the integral we used R’s `integrate()` function and to find the value of the integral equal to 1 we used the `uniroot()` function.

To find the spectral radius of the Poincaré map, 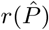, we used the method described in Liang et al. (2017, Lemma 2.5), which is similar to the Power method (Wikipedia, 2018), but specifically developed for delay differential equations. This algorithm involves numerically solving,

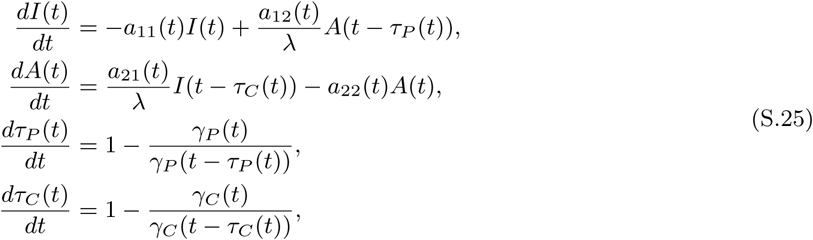

where 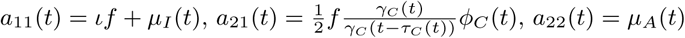, and 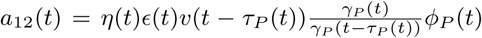.The system (S.21) is the same as the system of equations (1) from the main text except that 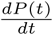 and 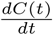 are removed because the two equations are decoupled from system (1) (see Section S.3), and the *a*_12_(*t*) and *a*_21_(*t*) terms are divided by *λ* to calculate *R*_0_ as detailed in equation (2.11) of Zhao (2017).

**Fig. S.3.**
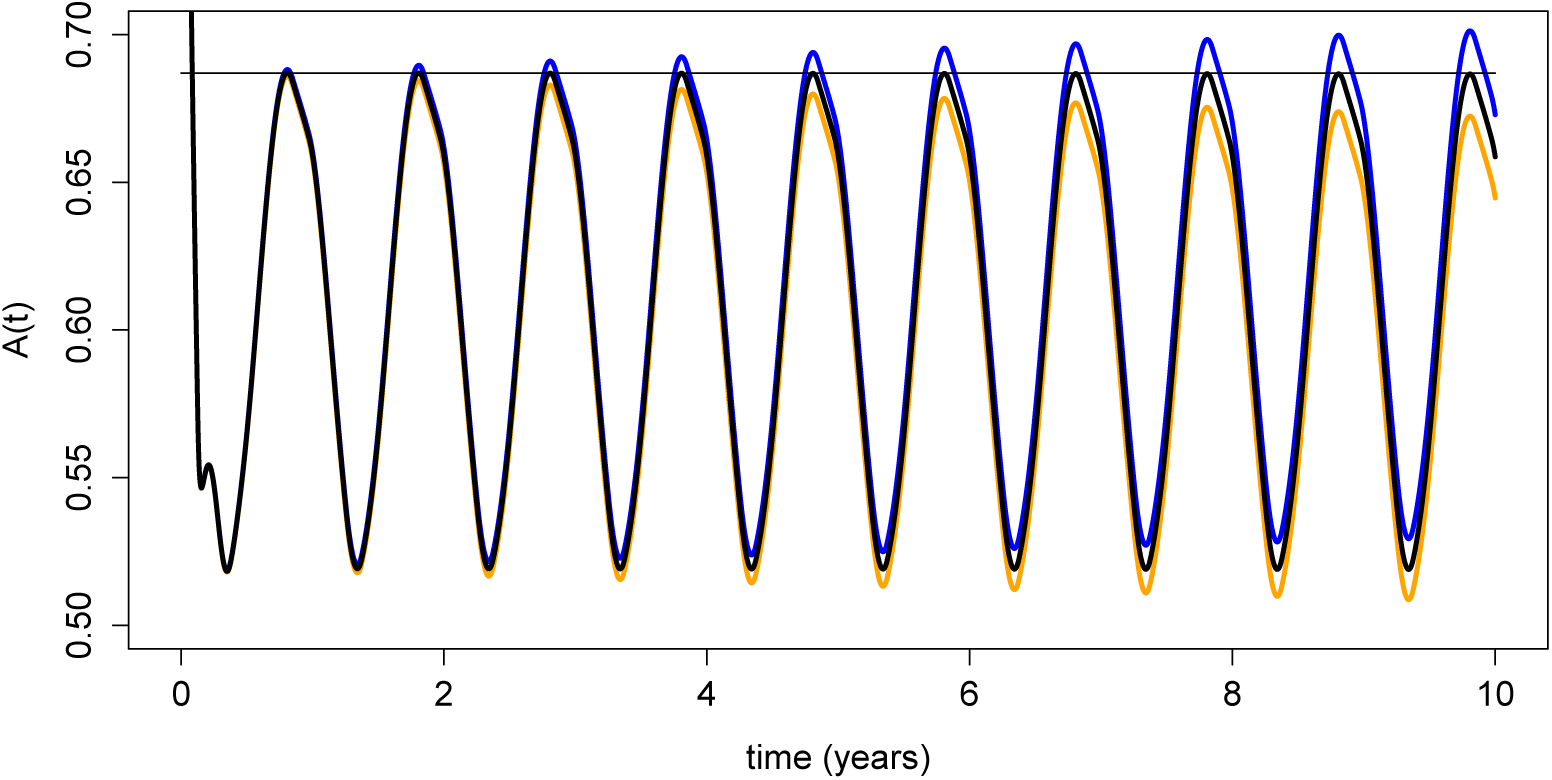
Numerical verification of *f*_*crit*_ as the threshold for population growth. Numerical solutions to the system (1) and (2) parameterized for the BCB site with *f* = *f*_*crit*_ (black), *f* = *f*_*crit*_−100 (orange), and *f* = *f*_*crit*_ + 100 (blue) confirm that *f*_*crit*_ is the threshold value of population growth. A horizontal line is shown at *A*(*t*) = 0.687 for reference.

The net reproductive ratio, *R*_0_, is the value of *λ* such that 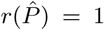, and we solved for this value using the `uniroot()` function. The Floquet exponent was calculated as *µ* = log(*M)/ω* when *λ* was set to 1. The critical stocking density was calculated as the value of *f* such that *µ* = 0 and was implemented using the `uniroot()` function. Figure S.3 shows the dynamics of (1) for *f*_*crit*_ (black), *f*_*crit*_ + 100 (blue), and *f*_*crit*_ − 100 (orange).

### S.5 The follow-up treatment window in seasonal environments

For a sea louse that became an adult female at time 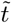, we back-calculate the time when the chalimus stage was reached, 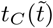, and when hatching occurred (i.e., the nauplius stage was first reached), 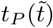, as:

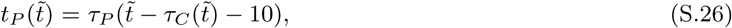

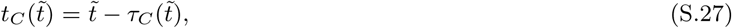

where 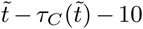 is the time when the copepodid stage was reached for a louse that became an adult female at 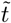. In Figure 4, we plot the time of an initial treatment, *t*_0_, on the x-axis. To plot the start of the follow-up treatment window in Figure 4, we identify salmon lice that hatched at the time of the initial treatment, 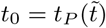, and plot the corresponding time to becoming a chalimus, 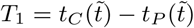 where the subtraction is to calculate the number of days to reach the chalimus stage, since 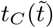 refers to time (i.e., 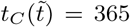 means the chalimus stage was reached on January 1st of year 2). To plot the end of the treatment window, we identify salmon lice that just became chalimi when the initial treatment occurred, 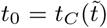 and plot the time to becoming an adult female,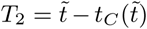. As for the constant temperature case, the treatment window exists for all sites we considered because *T*_1_ is always less than *T*_2_.

